# Cd-induced cytosolic proteome changes in the cyanobacterium *Anabaena* sp. PCC7120 are mediated by LexA as one of the regulatory proteins

**DOI:** 10.1101/2022.09.23.509143

**Authors:** Akanksha Srivastava, Arvind Kumar, Subhankar Biswas, Vaibhav Srivastava, Hema Rajaram, Yogesh Mishra

**Affiliations:** Department of Botany, Centre of Advanced Study in Botany, Institute of Science, Banaras Hindu University, Varanasi-221005, India; Molecular Biology Division, Bhabha Atomic Research Centre, Trombay, Mumbai-400085, India; Division of Glycoscience, Department of Chemistry, School of Engineering Sciences in Chemistry, Biotechnology and Health, Royal Institute of Technology (KTH), AlbaNova University Centre, Stockholm-10691, Sweden; Homi Bhabha National Institute, Anushakti Nagar, Mumbai-400094, India

**Author notes:** **Corresponding Authors:** Hema Rajaram; Yogesh Mishra.

**Keywords:** Electrophoretic mobility shift assay, LexA, Mass spectrometry, Real-time quantitative PCR, Transcription regulation, Two-dimensional gel electrophoresis

## Abstract

LexA, a well-characterized transcriptional repressor of the SOS genes in heterotrophic bacteria, has been shown to regulate diverse genes in cyanobacteria. An earlier study showed that LexA overexpression in a cyanobacterium, *Anabaena* sp. PCC7120 reduces its tolerance to Cd stress. This was later shown to be due to modulation of photosynthetic redox poising by LexA under Cd stress. However, in light of the global regulatory nature of LexA and the prior prediction of AnLexA-box in a few heavy metal-responsive genes, we speculated that LexA has a broad role in Cd stress tolerance, with regulation over a variety of Cd stress-responsive genes in addition to the regulation on genes related with photosynthetic redox poising. Thus, to further expand the knowledge on the regulatory role of LexA in Cd stress tolerance, a cytosolic proteome profiling of *Anabaena* constitutively overexpressing LexA upon Cd stress was performed. The proteomic study revealed 25 differentially accumulated proteins (DAPs) in response to the combined effect of LexA overexpression and Cd stress, and the other 11 DAPs exclusively in response to either LexA overexpression or Cd stress. The 36 identified proteins were related with a variety of functions, including photosynthesis, carbon metabolism, antioxidative defence, protein turnover, chaperones, post-transcriptional modifications, and a few unknown and hypothetical proteins. The regulation of LexA on corresponding genes, as well as six previously reported Cd efflux transporters, was further validated by the presence of AnLexA-boxes, transcript, and/or promoter analyses. In a nutshell, this study identifies the regulation of LexA on several genes and proteins of various functional categories in *Anabaena* that are responsive to Cd stress, hence expanding the regulatory role of LexA under Cd stress.

**Highlights:**

- LexA overexpression was earlier shown to decrease Cd stress tolerance in *Anabaena*.
- We examined the combined effect of LexA overexpression and Cd on *Anabaena* proteome.
- Upon LexA overexpression or Cd stress or both, 36 differential proteins were found.
- *In silico*, transcript and EMSA proved LexA regulation on them and few transporters.
- The findings of this study extended the regulatory role of LexA in Cd tolerance.

## 1. Introduction

LexA protein instead of being restricted to its role as an SOS response regulator as observed in *Escherichia coli* and other bacteria (Little et al., 1980: Wojciechowski et al., 1991; Clerch et al., 1996; Fernández de Henestrosa et al., 2000; Da Rocha et al., 2008), exhibits a global regulatory role in the cyanobacterium *Anabaena* sp. PCC7120 (Kumar et al., 2018). It was found to function both as a repressor and an activator across cyanobacterial species. In *Synechocystis* sp. PCC6803 (hereinafter referred to as *Synechocystis*), expression of *lexA* gene was not found to be inducible by DNA damage (Patterson-Fortin et al., 2006), and its depletion altered the expression of several genes, none of which belonged to the SOS-regulon, rather modulated C-starvation response (Domain et al., 2004; Kizawa et al., 2016). In *Synechocystis*, genes encoding bidirectional hydrogenase (Gutekunst et al., 2005), redox-sensitive RNA helicase (Patterson-Fortin et al., 2006), proteins involved in sodium-dependent bicarbonate transport (Lieman-Hurwitz et al., 2009), phototactic motility, accumulation of glucosylglycerol (Kizawa et al., 2016), fatty acid biosynthesis (Kizawa et al., 2017), and salt stress response (Takashima et al., 2020) were shown to be directly regulated by LexA. The lack of cleavable LexA in *Synechocystis* (Li et al., 2010) was speculated to be a reason for the DNA repair genes not being regulated by LexA. On the other hand, LexA of *Anabaena* sp. PCC7120 (hereinafter referred to as *Anabaena*), exhibited RecA-independent but pH-dependent cleavage (Kumar et al., 2015), and was found to function as a global regulator for genes belonging to several functional categories (Mazón et al., 2004; Sjöholm et al., 2007; Kirti et al., 2017; Kumar et al., 2018; Pandey et al., 2018; Rajaram et al., 2022; Pradhan et al., 2022 PREPRINT; Srivastava et al., 2022a; Srivastava et al., 2022b PREPRINT). The regulation by LexA in *Anabaena* was through binding to the AnLexA-Box defined as AGT-N_4-11_-ACT with one base mismatch allowed within a 4-base palindrome (Kumar et al., 2018; Srivastava et al., 2022a). The binding has been shown for a few DNA repair genes (Kirti et al., 2017; Kumar et al., 2018; Pandey et al., 2018; Pradhan et al., 2022 PREPRINT), oxidative stress-alleviating and C-metabolism genes (Kumar et al., 2018), photosynthetic electron transport chain (pETC) components (Srivastava et al., 2022a), and a few γ-radiation response genes (Srivastava et al., 2022b PREPRINT). The global regulatory role of LexA, both as an activator and a repressor, may be the reason for the non-viable nature of the *lexA* mutant of *Anabaena* (Kumar et al., 2018).

Cyanobacteria exhibit a remarkable ability to survive cadmium (Cd) stress by inducing changes at morpho-ultrastructural, biochemical, physiological, and molecular levels (Jensen et al., 1982; Blasi et al., 2012; Singh et al., 2015; Victoria et al., 2018; Ahad and Syiem, 2021; Shen et al., 2021; Srivastava et al., 2021a,b; Tian et al., 2022). Our earlier results showed decreased Cd tolerance in *Anabaena* upon overexpression of LexA (Kumar et al., 2018), which was later shown to be through the modulation of photosynthetic redox poise as one of the regulatory components (Srivastava et al., 2022a). Since some of the heavy metal-responsive genes were shown to possess the AnLexA-Box (Kumar et al., 2018), we decided to expand further our knowledge on the involvement of LexA in Cd tolerance by conducting a proteomic study in *Anabaena* cells constitutively expressing LexA upon exposure to Cd stress, followed by validation of expression through real-time quantitative PCR (RT-qPCR) and electrophoretic mobility shift assay (EMSA) studies with LexA. The findings of the study showed that, in addition to regulating photosynthetic genes and modulating photosynthetic redox poising under Cd stress, LexA regulates the expression of several genes from various functional categories that respond to Cd stress and help in Cd efflux as well as the induction of cellular defence and repair systems.

## 2. Material and methods

### 2.1. Strains used, growth, and stress conditions

The recombinant *Anabaena* strains, vector control AnpAM (Rajaram & Apte, 2010) and LexA-overexpressing An*lexA*^+^ (Kumar et al., 2018) were grown at 25 ± 2 °C under continuous illumination of 72 μmol photon m^−2^ s^−1^ photosynthetic photon flux density in nitrogen-supplemented BG-11 medium, pH 7.2 (Castenholz, 1988) with 10 μg neomycin mL^−1^. Cd stress treatment was performed by concentrating 5-day-old recombinant *Anabaena* cells to 10 μg Chl *a* mL^−1^ and exposing them to 10 μM CdCl_2_ for 1 day, as described earlier (Srivastava et al., 2021a).

### 2.2 Protein extraction and two-dimensional (2-D) gel electrophoresis

Proteins from recombinant *Anabaena* cultures were extracted according to Kaur et al. (2019), precipitated overnight at −20 °C with six volumes of ice-chilled 10% TCA-acetone, and resuspended in a solubilization buffer containing 7 M urea, 2 M thiourea, 4% CHAPS, 40 mM DTT, and 1.0 % IPG buffer. Protein estimation was carried out by using the Bradford method (Bradford, 1976) and electrophoretically separated by 2-D gel electrophoresis, as described earlier (Srivastava et al., 2021b). Excision and identification of differentially accumulated protein (DAP) spots were carried out as described in detail earlier (Srivastava et al., 2021a,b). The peptides were selected based on mascot scores that met or exceeded the statistical significance threshold (*p* < 0.05).

### 2.3 In silico prediction of the promoter region and AnLexA-box

The upstream region (300 – 500 bases) of specific genes was analyzed for probable bacterial promoters using BPROM software (http://www.softberry.com/berry.phtml?topic) (Solovyev & Salamov, 2011) and AnLexA-box as defined earlier (Kumar et al., 2018; Srivastava et al., 2022a).

### 2.4 Electrophoretic mobility shift assay (EMSA)

EMSA studies were carried out using 200 – 300 bp DNA fragments corresponding to upstream regions of genes as listed in supplementary Table S1 incubated with varying concentrations of LexA, as described previously (Kirti et al., 2017). SYBR Dye I was used to stain the gels, which were then viewed with a UV trans-illuminator and densitometrically evaluated with ImageJ software (Schneider et al., 2012).

### 2.5 Transcript analysis

Total RNA from recombinant *Anabaena* strains was extracted, and RT-qPCR analysis was carried out using specific primer pairs (Supplementary Table S2), as described earlier (Srivastava et al., 2021a,b). Two housekeeping genes, *rnpA* and *secA*, were used to normalize the results (Pinto et al., 2012). The relative transcript level was estimated using the 2^−ΔΔCt^ method (Livak and Schnittgen, 2001). A significant difference in mRNA levels was considered only when the fold change was ≥ 1.3 (Student’s t-Test *p* < 0.05).

## 3. Results

Since overexpression of LexA in *Anabaena* was shown to decrease tolerance to Cd stress (Kumar et al., 2018), it was postulated that other than the effect on photosynthetic redox poising during Cd stress as shown earlier (Srivastava et al., 2022a), LexA as a regulator could affect the expression of genes involved in metal export or proteins involved in mitigating effect of Cd stress. In this direction, the combined effect of Cd stress and LexA overexpression on *Anabaena* proteome was analyzed, followed by the validation of the observed proteomic results in terms of the role of LexA as a regulator by qRT-PCR and EMSA.

### 3.1 Effect of LexA on modulation of Cd stress-responsive cytosolic proteins in Anabaena

The proteome profile of unstressed cultures of AnpAM and An*lexA*^+^ has been evaluated earlier (Kumar et al., 2018; Srivastava et al., 2022b PREPRINT), and also that of wild-type *Anabaena* in response to Cd stress (Singh et al., 2015). Thus, here we have compared the 2-D proteome profile between the unstressed (US) culture of AnpAM (Fig. 1A) and the Cd-stressed (CdS) culture of An*lexA*^+^ (Fig. 1B) to assess the response of Cd stress in the presence of overexpressed LexA protein. Approximately 250 spots, in the molecular weight range of 8–240 kDa and pI range of 3–10, were detected on each gel (Fig. 1), of which about 170 protein spots could be reliably matched on both the gels and their triplicates. Of these, the accumulation of 28 protein spots differed by at least 1.5-fold (Student’s t-Test *p* < 0.05) (Fig. 1 and Table 1). In addition to these, 11 spots exhibited at least a 1.5-fold change in response to either Cd stress in AnpAM cells or LexA overexpression, as shown in Fig. S1 and tabulated in Table S3. All 39 DAP spots were identified using reversed-phase LC-ESI-MS/MS, and their fold changes and identification details are provided in Tables 1 and S3.

**Fig. 1.**
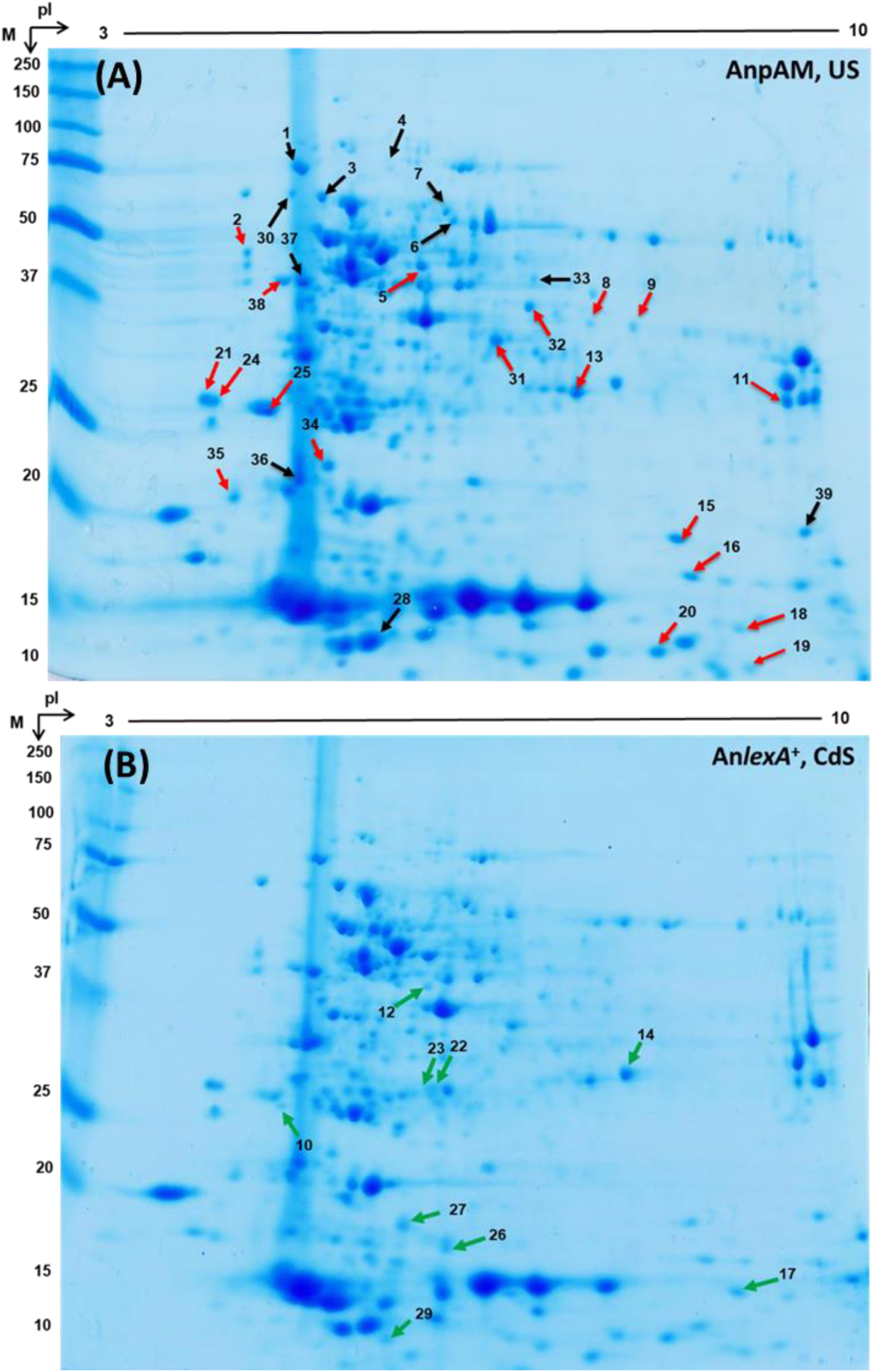
Two-dimensional (2-D) gel electrophoresis-based cytosolic proteome analysis of *Anabaena* sp. PCC7120 in response to Cd stress in the presence of overexpressed LexA protein. The recombinant *Anabaena* strains (A) AnpAM and (B) An*lexA*^+^, respectively, stand for vector control and LexA-overexpressing strains. US and CdS respectively, indicate unstressed and Cd-stressed (CdS) cultures. Differentially accumulated cytosolic proteins are labelled with their corresponding spot numbers with red and green arrows. Red arrows in (A) or green arrows in (B), respectively, represent protein spots that showed lower or higher accumulation in response to Cd stress in the presence of constitutively expressed LexA protein. The molecular masses (kDa) of protein size markers are displayed on the left, while the pI (3-10) is presented on the top.

**Table 1.** Identification details, as well as mean fold changes in the intensities of differentially accumulated protein (DAP) spots in *Anabaena* sp. PCC7120 in response to Cd stress in the presence of overexpressed LexA protein

| Spot No. | Gene No. | Protein name<sup>#</sup> | Functional category (FC) | Swiss Prot No | Molecular mass (kDa) and pI | | Fold change in An$\text{lexA}^+$, CdS compared to AnpAM, US |
| --- | --- | --- | --- | --- | --- | --- | --- |
|  |  |  |  |  | Observed | Predicted |  |
| 2 | *all4337* | Elongation factor Tu (EF-Tu) | III | Q8YP63 | 46.55/4.20 | 44.84/5.25 | **-2.63** |
| 5 | *alr0803* | Alr0803 (Hypo) | V | Q8YYP3 | 44.83/5.76 | 43.57/5.61 | **-2.00** |
| 8 | *all0865* | Carboxysome assembly protein (CcmM) | I | Q8YYI3 | 36.21/7.11 | 59.82/6.77 | **-2.68** |
| 9 | *all0865* |  |  |  | 36.13/7.32 |  | **-3.03** |
| 10 | *all1864* | All1864 (~NADH-DH) | V | Q8YVV9 | 24.74/4.39 | 27.14/4.75 | **2.58** |
| 11 | *alr0535* | Phycobilisome rod-core linker polypep. CpcG2 | I | P29987 | 27.05/9.11 | 28.63/9.06 | **-2.27** |
| 12 | *all1865* | All1865 (~NemA) | V | Q8YVV8 | 37.33/5.45 | 43.87/5.49 | **5.72** |
| 13 | *alr0882* | Alr0882 (Hypo) | V | Q8YYG8 | 29.17/7.11 | 30.87/6.84 | **-2.88** |
| 14 | *all2425* | All2425 (Unknown) | V | Q8YUC7 | 30.43/7.33 | 28.02/6.90 | **1.82** |
| 15 | *alr1208* | Ribosome-recycling factor (RRF) | III | Q8YXK4 | 18.78/8.11 | 20.22/7.82 | **-4.14** |
| 16 | *all0579* | 50S ribosomal protein L9 (Rpl9) | III | Q8YZA0 | 16.71/8.18 | 16.75/7.93 | **-2.46** |
| 17 | *all7597* | All7597 (Hypo) | V | Q8ZSB4 | 14.98/8.71 | 16.90/9.07 | **8.33** |
| 18 | *all0329* | Photosystem I reaction center subunit II (PsaD) | I | P58573 | 11.05/8.89 | 15.14/9.30 | **-4.85** |
| 19 | *all0329* |  |  |  | 13.57/8.95 |  | **-1.69** |
| 20 | *all0258* | Plastocyanin (PetE) | I | P46444 | 12.53/8.52 | 14.66/9.10 | **-1.82** |
| 21 | *alr0806* | Alr0806 (Unknown) | V | Q8YYP0 | 24.87/4.12 | 17.68/4.42 | **-2.55** |
| 24 | *alr0806* |  |  |  | 24.83/4.15 |  | **-2.43** |
| 22* | *all4391* | Enoyl-[acyl-carrier-protein] reductase [NADH] (ENR) | IV | Q05069 | 26.93/5.90 | 27.52/5.56 | **>10.00** |
| 23* | *alr0782* | Ribulose-phosphate 3-epimerase (Rpe) | I | Q8YYR4 | 26.14/5.56 | 25.74/5.39 | **>10.00** |
| 25 | *alr4688* | All4688 (Hypo) | V | Q8YN82 | 25.78/4.45 | 32.04/4.63 | **-4.76** |
| 26 | *alr5302* | 50S ribosomal protein L10 (Rpl10) | III | Q8YLJ6 | 16.71/5.84 | 19.48/5.71 | **1.73** |
| 27 | *all4894* | All4894 (Hypo) | V | Q8YMN9 | 16.98/5.61 | 14.57/6.71 | **1.92** |
| 29 | *alr2205* | Thioredoxin (TrxO) | II | Q8YUX6 | 10.57/5.21 | 12.91/5.28 | **6.48** |
| 31 | *all5062* | Gly-3-phosphate<br>DH 2 (Gap-2) | I | P58554 | 34.87/6.12 | 37.12/6.09 | **-2.51** |
| 32 | *all3909* | Uroporphyrinogen<br>decarboxylase<br>(HemE) | I | Q8YQC4 | 36.83/6.81 | 38.79/6.33 | **-4.37** |
| 34 | *all3797* | All3797 (Hypo) | V | Q8YQM3 | 22.14/4.85 | 27.87/7.90 | **-2.5** |
| 35 | *all2110* | All2110 (Hypo) | V | Q8YV71 | 21.48/4.25 | 22.27/4.45 | **-2.55** |
| 38 | *all0136* | 30S ribosomal<br>protein S1 (Rps1) | III | Q8Z0G0 | 38.11/4.35 | 38.34/4.68 | **-1.66** |

In response to the combined effects of Cd stress and LexA overexpression, 19 DAP spots (2, 5, 8, 9, 11, 13, 15, 16, 18, 19, 20, 21, 24, 25, 31, 32, 34, 35, and 38) showed decreased accumulation and 9 DAP spots (10, 12, 14, 17, 22, 23, 26, 27, and 29) showed increased accumulation when compared with AnpAM, US (Fig. 1 and Table 1). Of the remaining identified DAPs, (i) 4 DAP spots (6, 7, 36, and 39) exhibited decreased accumulation when the cells were either exposed to Cd stress or the cells overexpressed LexA, but not both (Fig. S1 and Tables S3), (ii) spot no. 1 (DnaK) showed decreased accumulation and spot no. 28 (TrxA) increased accumulation only upon LexA overexpression (Fig. S1 and Table S3), conforming to the earlier results (Kumar et al., 2018), and (iii) 5 DAPs (3, 4, 30, 33, and 37) showed increased accumulation under Cd stress but decreased accumulation upon LexA overexpression (Fig, S1 and Table S3). These data indicated the involvement of LexA in regulating Cd responses of *Anabaena*.

Principal component analysis (PCA) of AnpAM, US; AnpAM, CdS; An*lexA*^+^, US; and An*lexA*^+^, CdS using BioDiversity Pro 2.0 (McAleece et al., 1997) revealed clear separation among the four samples (Fig. 2A), indicating a significant difference in their cytosolic proteomes. This data also indicates the substantial involvement of LexA in regulating Cd stress-induced cytosolic proteomic changes in *Anabaena*. Few protein spots were segregated from others in the PCA of 39 DAP spots (Fig. 2B); therefore, they were considered probable outliers and given priority for mass spectrometry investigation.

**Fig. 2.**
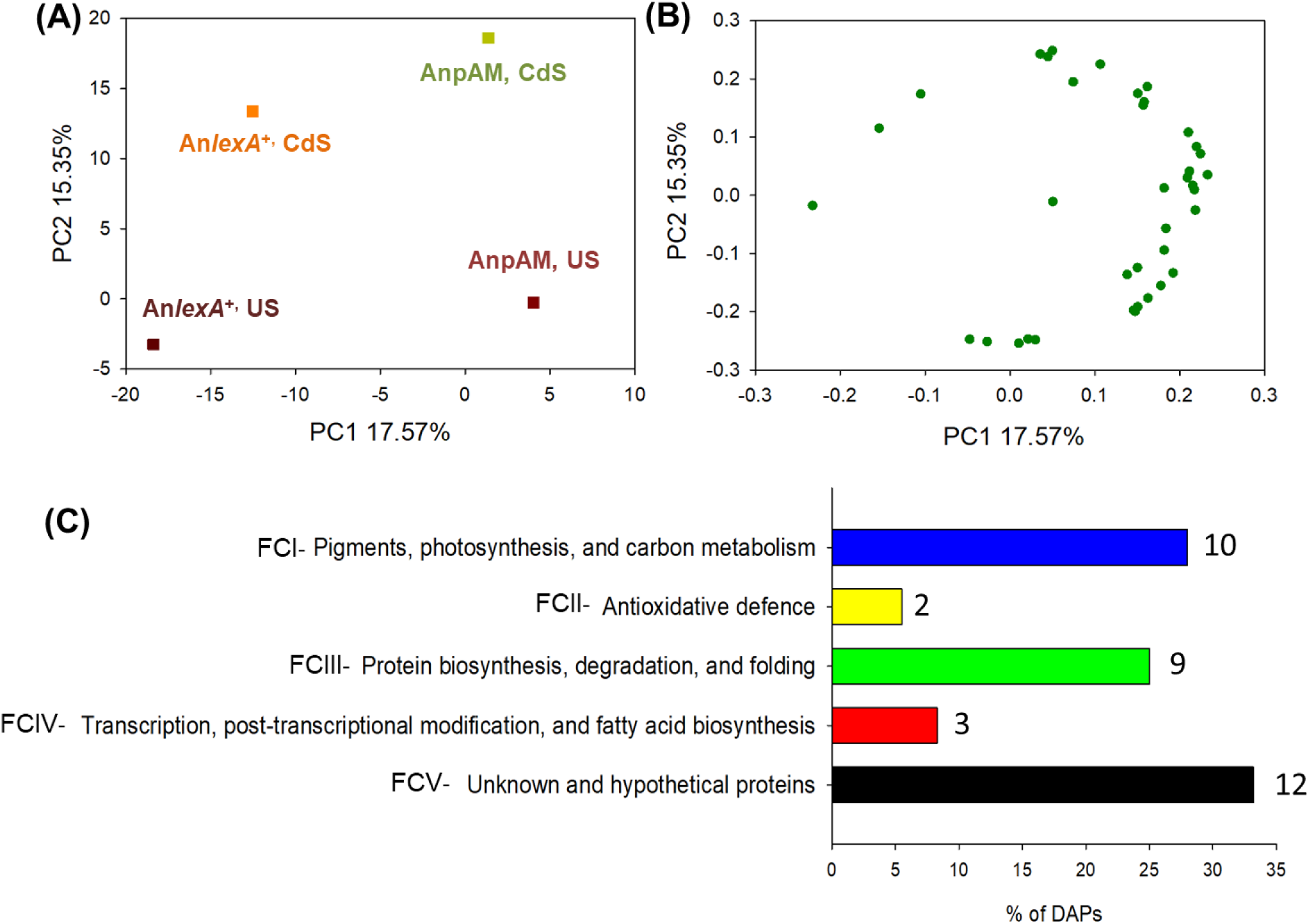
Principal component analysis (PCA) of 2D-PAGE data using BioDiversity Pro 2.0 (A) PCA was performed on the cytosolic proteomes of two recombinant *Anabaena* strains, vector control (AnpAM) and LexA-overexpressing, (An*lexA*^+^) under unstressed (US) and Cd-stressed (CdS) conditions. (B) PCA of 39 DAP spots identified in response to Cd stress and LexA overexpression either individually or combined (C) Classification of 36 unique proteins identified from 39 DAP spots into five (I-V) functional categories (FCs). The number of different proteins that each FC has is indicated near its bar.

The identification of proteins indicated that 39 DAP spots corresponded to 36 different proteins (Table 1, S3), with three proteins (CcmM, PsaD, and Alr0806) identified in two spots each. Of these, 16 proteins (EF-Tu, Alr0803, CcmM, CpcG2, Alr0882, RRF, Rpl9, PsaD, PetE, Alr0806, All4688, Gap-2, HemE, All3797, All2110, and Rps1), showed decreased abundance when *Anabaena* cells overexpressing LexA (An*lexA*^+^) were subjected to Cd stress (Cds) compared to unstressed vector control *Anabaena* cells (AnpAM, US) (Fig. 1 and Table 1). Under these conditions, abundance of 9 proteins (All1864, All1865, All2425, All7597, ENR, Rpe, Rpl10, All4894, and TrxO) was found to increase (Fig. 1 and Table 1). Among the DAPs showing a change in abundance in the presence of only one of the two conditions, i.e. either Cd stress or LexA overexpression, each of them corresponded to a different protein, resulting in the identification of 11 proteins, namely DnaK2, AtpA, OpA, ICDH, PGDH, TrxA, NusA, AMT, CpcB, All4050, and Rbp (Fig. S1 and Table S3). During this study, we identified a few additional proteins, which were not previously reported to respond under Cd stress (Singh et al., 2015) or LexA overexpression (Kumar et al., 2018; Srivastava et al., 2022b PREPRINT). The functional classification of 36 identified DAPs in response to either Cd stress or LexA overexpression or both indicated that LexA primarily modulates the accumulation of photosynthesis, carbon metabolism, protein turnover, and a substantial number of unknown and hypothetical proteins (Fig. 2C and Table 1, S3).

### 3.2 Assessment of LexA-mediated transcriptional regulation of genes encoding the identified cytosolic DAPs, as well as a few Cd efflux transporters

As seen in Tables 1 and S3, all the identified DAPs exhibited differential accumulation upon LexA overexpression irrespective of the absence or presence of Cd stress. To assess if this differential accumulation is indeed regulated through LexA, they were examined for the (i) presence of AnLexA-boxes in their upstream regulatory regions, (ii) EMSA studies to assess LexA binding to the predicted AnLexA-boxes, and (iii) RT-qPCR analysis for nine selected genes in response to Cd or LexA overexpression or both. Since membrane proteins are difficult to separate on 2-D PAGE (Wilkins et al., 1998) and were excluded in our study of the cytosolic proteome, the genes encoding a few previously identified Cd efflux transporters, such as *all7618*, *all7619*, *ycf16*, *aztA*, *alr4267*, and *secD* (Xu et al., 2018; Victoria et al., 2018; Srivastava et al., 2021b), were also included in this study to assess if LexA could be involved in regulating the expression of efflux proteins as well.

*In silico* prediction revealed that out of 36 DAPs encoding genes, 32 have one or more AnLexA-boxes in their upstream regions, with two of them, *alr1208* and *alr4853,* having the promoter and regulatory regions upstream of the gene preceding it (Table 2). Two more of them, *all1864* and *all1865*, are part of a tricistronic operon, with the regulatory elements present upstream of the first gene of the operon, *all1866*, encoding Thioredoxin (Table 2). The two genes predicted not to be regulated by LexA are *tufA* and *atpA,* which lack the AnLexA-Box and two genes *alr0803* and *petE*, wherein the AnLexA-Box is distant from the promoter region (Table 2). Among the six metal transporters analyzed, all except *secD* were predicted to have a regulatory AnLexA-Box (Table 2).

**Table 2.**
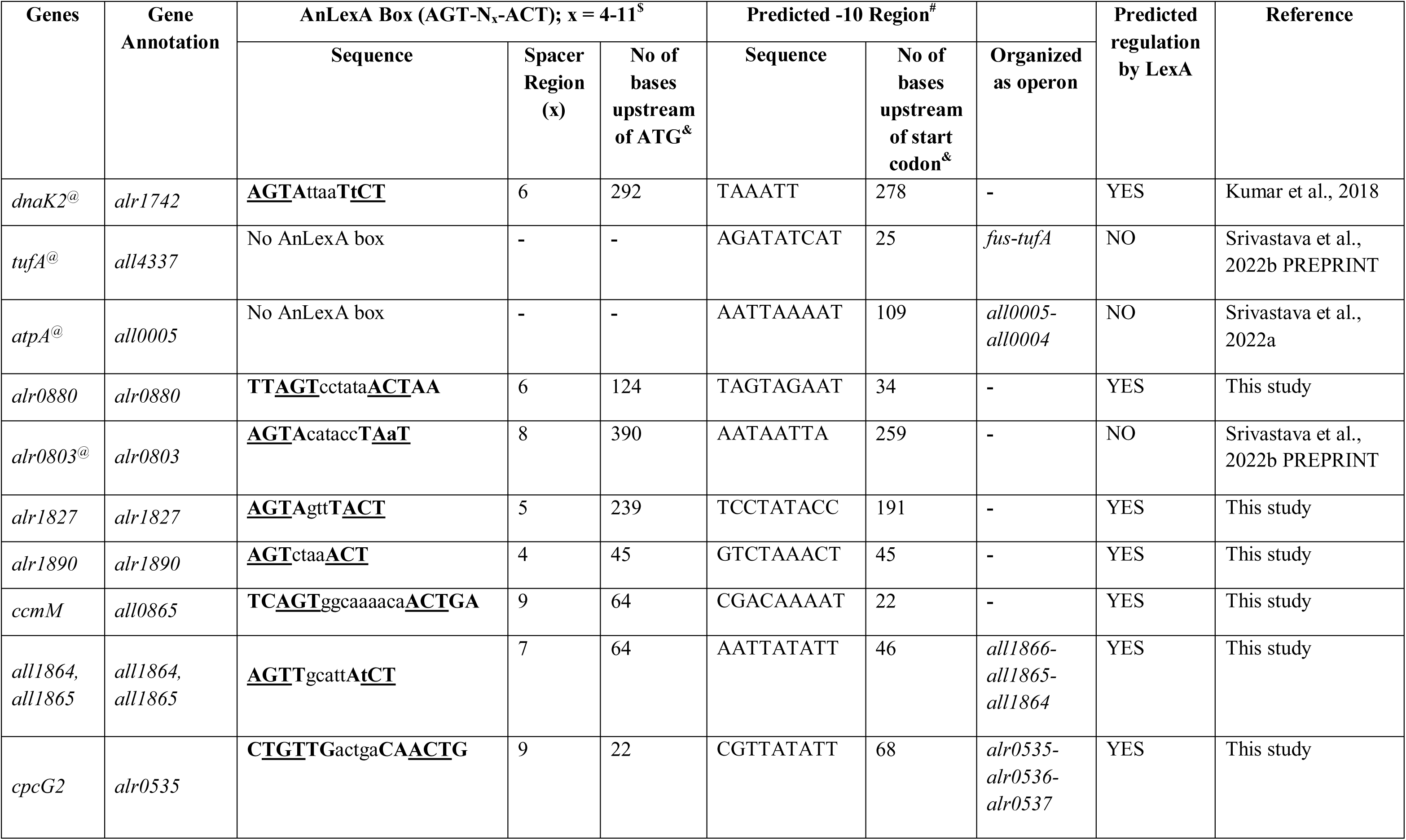
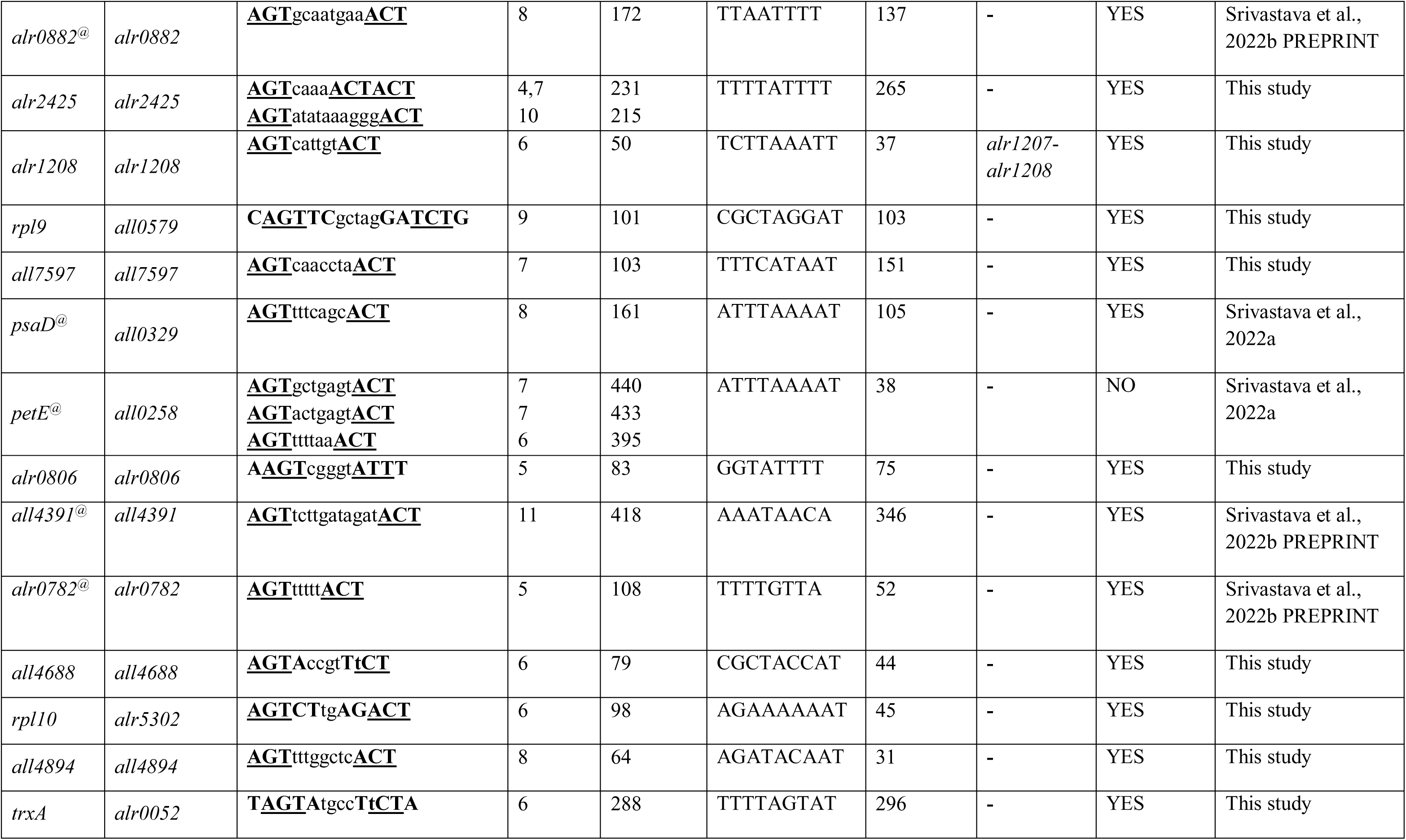
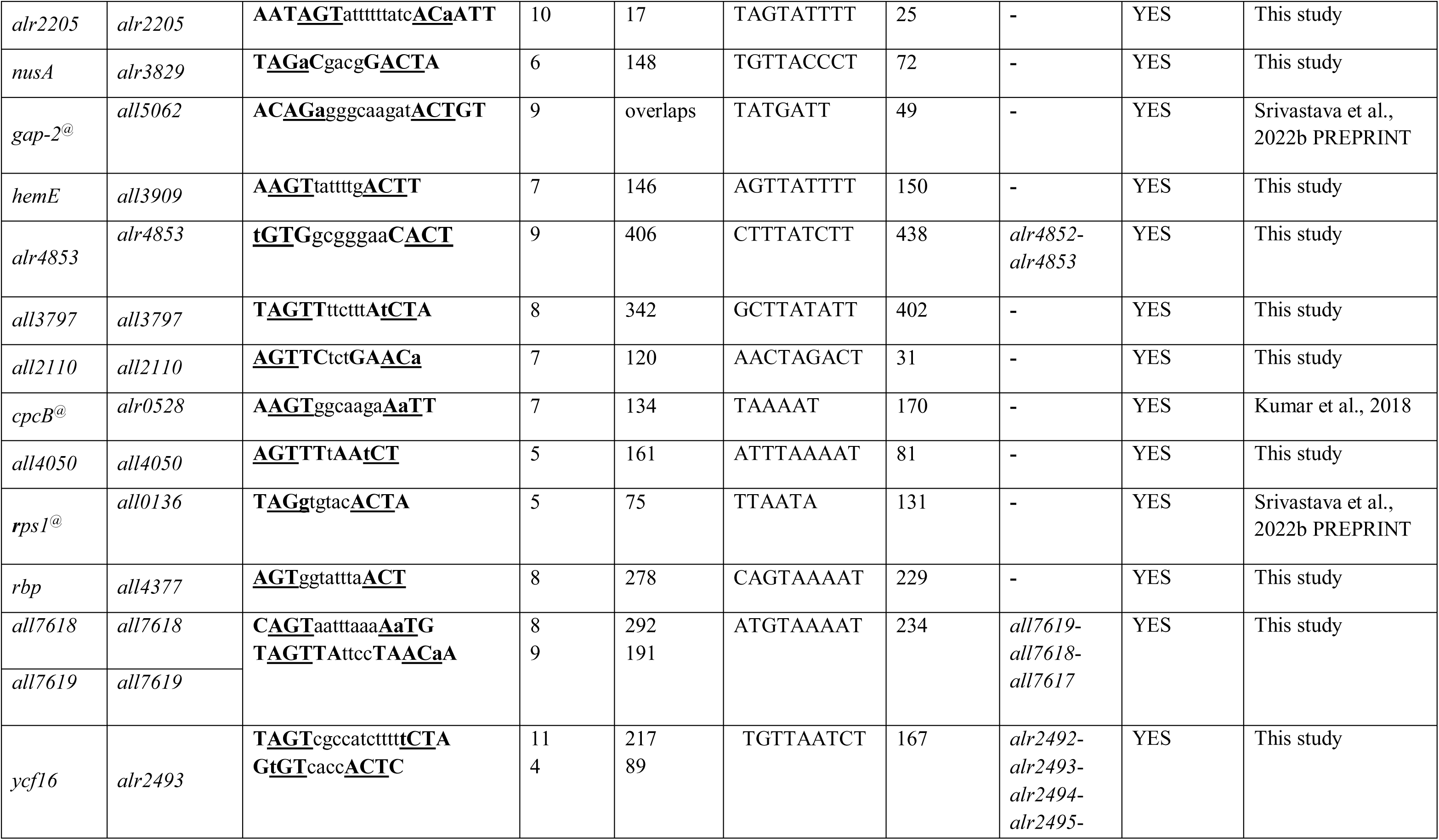
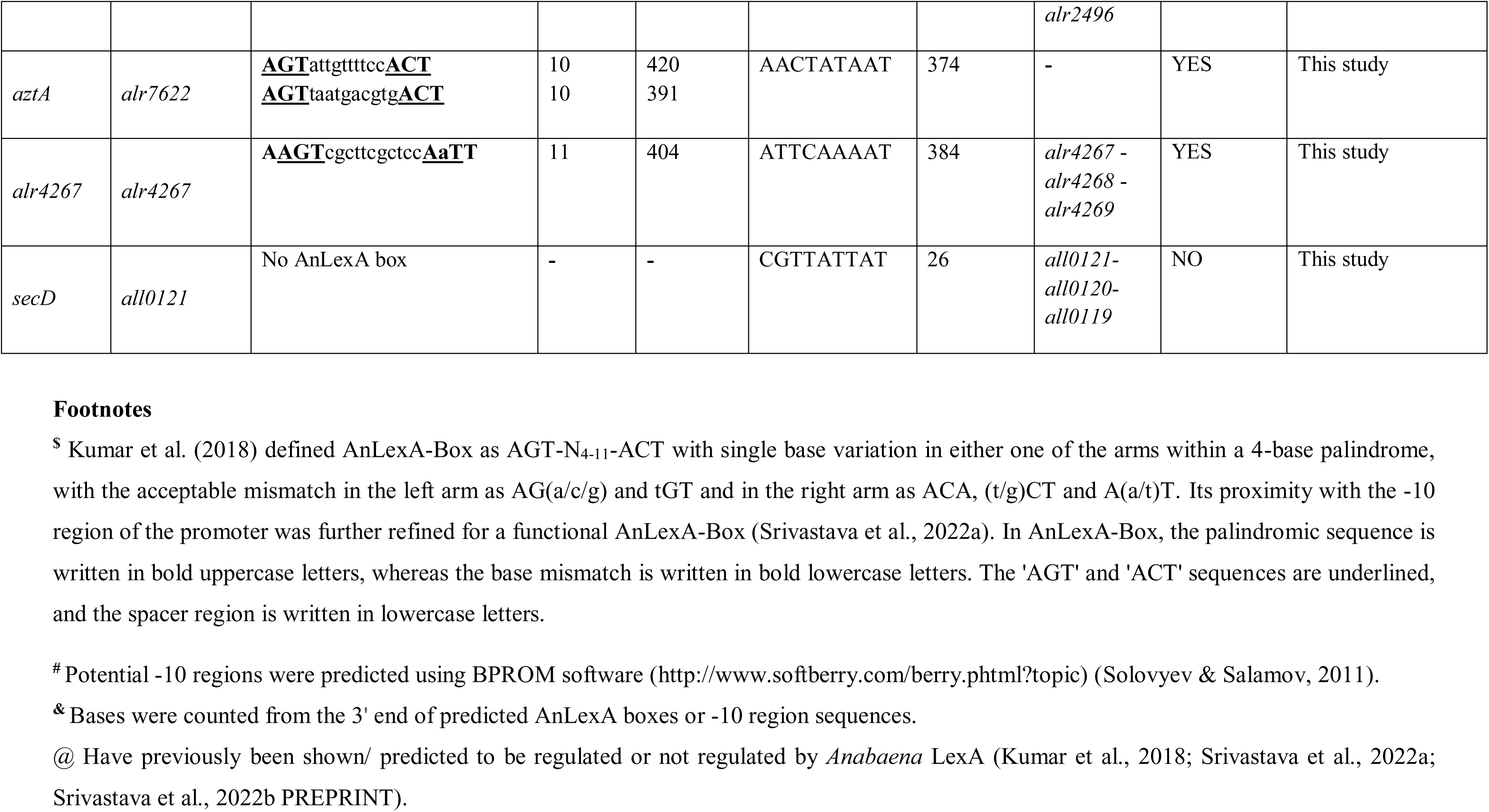
*In silico* AnLexA-Box prediction in the upstream regulatory regions of genes coding for identified cytosolic DAPs, as well as Cd efflux transporters

Of 32 DAPs encoding genes with potential AnLexA-box in their promoter regions, four genes, namely *dnaK2*, *psaD*, *gap-2*, and *cpcB* have previously been experimentally demonstrated to be regulated by *Anabaena* LexA (Kumar et al., 2018; Srivastava et al., 2022a; Srivastava et al., 2022b PREPRINT), and four additional genes, namely *alr0882*, *all4391*, *alr0782*, and *rps1* have previously been predicted to be regulated by *Anabaena* LexA (Srivastava et al., 2022b PREPRINT). Furthermore, the *in vitro* interaction between purified *Anabaena* LexA and upstream region of 5 genes encoding identified DAPs (*ccmM*, *all1864*, *alr2205*, *nusA*, and *hemE*) and 3 Cd efflux transporters (*all7619*, *ycf16*, and *aztA*) was examined (Fig. 3). All tested DNA fragments exhibited 50–70% binding even at 62 nM LexA; however no binding was seen for the negative control DNA fragment lacking the AnLexA-Box (Fig. 3).

**Fig. 3.**
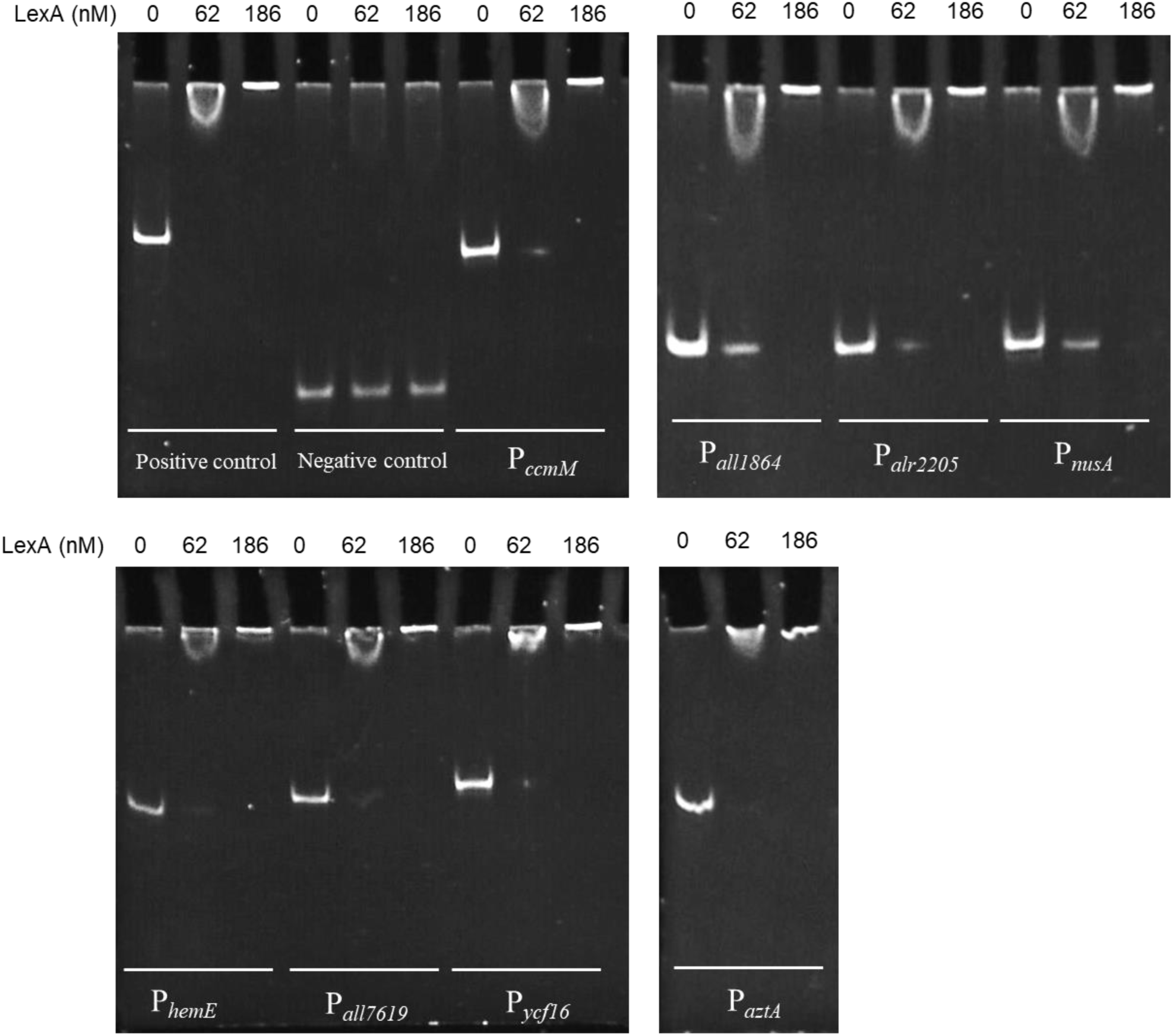
Binding of *Anabaena* LexA to the upstream region of 5 genes encoding identified DAPs (*ccmM*, *all1864*, *alr2205*, *nusA*, and *hemE*) and 3 Cd efflux transporters (*all7619*, *ycf16*, and *aztA*) assessed by Electrophoretic mobility shift assay (EMSA). Different concentrations of purified *Anabaena* LexA (0, 62 and/or 186 nM) were incubated with amplified promoter fragments (∼30 ng) as indicated, followed by electrophoretic separation on 8% non-denaturing polyacrylamide gel, staining with SYBR Green I, and visualisation using UV trans-illuminator. The *lexA* promoter region was used as a positive control, while the promoter region of a gene lacking the AnLexA-Box was used as a negative control.

Transcript analysis of selected genes among the identified DAPs revealed increased transcription of *alr0880*, *all1864*, *alr2205*, and *all2110*; decreased transcription of *ccmM*, *rpl9*, and *hemE*, and no change in that of *nusA* and *all4050* in combined presence of LexA and Cd [An*lexA*^+^, CdS vs AnpAM, US] (Fig. 4A). The data agreed with the observed changes in proteomics data with the exception of Alr0880 and All2110 (Table 1, S3), Of these genes, overexpression of LexA under unstressed conditions [An*lexA*^+^, US vs AnpAM, US] resulted in up-regulation of transcription of *ccmM*, *nusA*, and *all4050*, while that of *all2110* decreased, and the remaining 5 genes showing no discernible change (Fig. 4B). On the other hand, Cd stress on vector control cells [AnpAM, CdS vs AnpAM, US] resulted in increased transcription of *alr0880*, *all1864*, *alr2205*, and *nusA* and decreased transcription of *ccmM*, *rpl9*, *hemE*, and *all4050*, while *all2110* exhibited no change in transcript levels (Fig. 4C). Comparison of the transcript expression data under three conditions, revealed that the genes showing differential expression under Cd stress in the presence of LexA, exhibited differential expression in the presence of at least one of the two individual conditions (only Cd stress or only LexA overexpression). The opposite responses of transcriptional regulation of *nusA* and *all4050* under LexA overexpression (Fig. 4B) and Cd stress (Fig. 4C) possibly resulted in no change detected under combined Cd stress and LexA overexpression (Fig. 4A). Transcript analysis was also carried out for 6 transporter genes, namely *czcA*, *czcB*, *ycf16*, *aztA*, *alr2267*, and *secD*. Exposure to Cd stress in LexA overexpressing cells, An*lexA*^+^ resulted in increased expression of *ycf16* and *secD*, while that of others remained unchanged (Fig. 4D). On the other hand, LexA overexpression increased the transcription of all 6 genes, including *secD* (Fig. 4E). Absence of AnLexA-box upstream of *secD* suggests indirect regulation by LexA through other LexA-regulated transcriptional factors. Cd stress, *per se*, i.e. in AnpAM cells, enhanced the expression of *ycf16* and *secD*, while down-regulating that of *czcA*, *czcB* and *alr4267* (Fig. 4F).

**Fig. 4.**
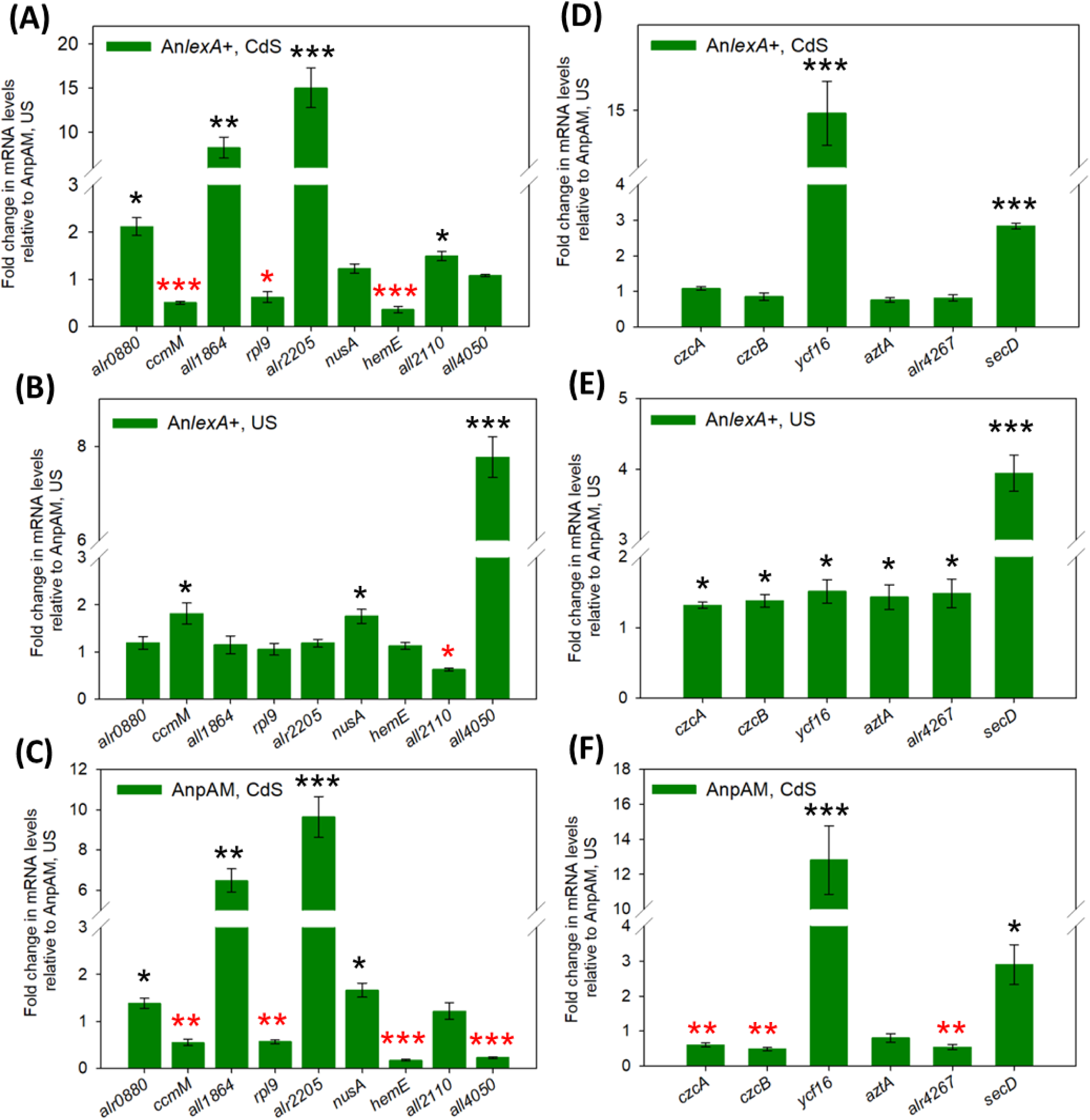
Transcript analysis of genes coding for nine selected cytosolic DAPs identified from 2D-PAGE namely, Alr0880, CcmM, All1864, Rpl9, TrxO (Alr2205), NusA, HemE, All2110, and All4050 (A, B, and C), as well as with six previously reported Cd efflux transporters, All7618, All7619, Ycf16, AztA, Alr4267, and SecD (D, E, and F) in two recombinant *Anabaena* strains, vector control (AnpAM) and LexA-overexpressing, (An*lexA*^+^) under unstressed (US) and Cd-stressed (CdS) conditions using real-time quantitative PCR (RT-qPCR). Transcript data from An*lexA*^+^, CdS were normalized to AnpAM, US to determine the combined effect of Cd stress and LexA overexpression (A and D). While, to assess the individual effects of LexA overexpression (B and E) and Cd stress (C and F), respectively, transcript data from An*lexA*^+^, US and AnpAM, CdS were each normalized to AnpAM, US. The fold change in transcript level was only considered significant when it exceeded the threshold value of 1.3 fold and also satisfied *p* < 0.05). Based on Student’s t-test with Tukey’s post-hoc comparison test, up-regulated genes are shown with black asterisk, whereas down-regulated genes are shown with red asterisk. Single asterisk (*) represents *p* < 0.05, double asterisk (**) represents *p* < 0.01, and triple asterisk (***) *p* < 0.001.

**Fig. 5.**
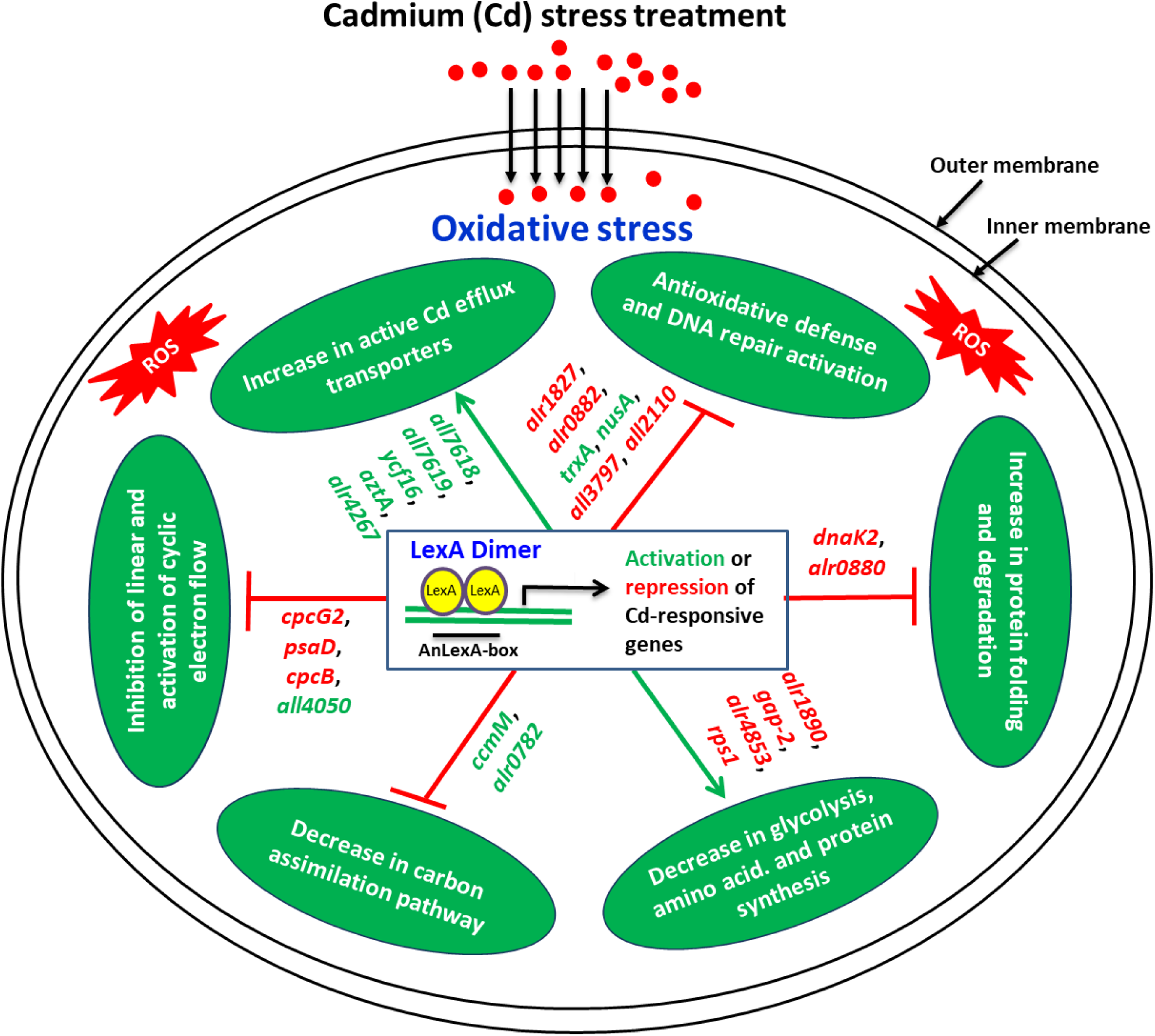
Schematic diagram showing the regulation of LexA protein on several Cd-responsive genes, as well as Cd stress response of *Anabaena* sp. PCC7120. Positively regulated Cd-responsive genes by LexA are highlighted in green, whereas negatively regulated genes are highlighted in red. The proteins encoded by these genes (symbols are italicized) are shown in Table 1. The Cd stress-induced adaptive responses of *Anabaena* are mentioned in white in green boxes as per the findings of this study, as well as Singh et al. (2015) and Srivastava et al. (2021a) and the possible regulation of LexA on these responses through activation or repression of different Cd-responsive genes is shown by a green arrow (→) for activated responses and a red inhibitory arrow (T) for repressed responses.

## 4. Discussion

Overexpression of LexA was previously shown to reduce Cd stress tolerance in *Anabaena* (Kumar et al., 2018), and later it was shown to be due to the modulation of photosynthetic redox poising (Srivastava et al., 2022a). However, due to the global regulatory nature of LexA in *Anabaena* and presence of AnLexA-box upstream of a few heavy metal-responsive genes (Kumar et al., 2018), we investigated the regulatory role of LexA on expression of Cd-responsive and Cd efflux genes, using the approaches of cytosolic proteome study followed by validation through RT-qPCR and EMSA in *Anabaena* cells overexpressing LexA and subjected to Cd stress.

### 4.1 Overexpression of LexA modulated the expression of photosynthetic, oxidative stress alleviation, protein turnover, and other Cd stress-responsive genes in Anabaena

From proteome analysis, we identified 25 distinct DAPs (corresponding to 28 spots) in Cd-stressed An*lexA*^+^ cells compared to unstressed AnpAM cells (Fig. 1 and Table 1), and 11 distinct DAPs, which were observed in either unstressed An*lexA*^+^ or Cd-stressed AnpAM cells compared to unstressed AnpAM cells (Fig. S1 and Table S3). These proteins were then categorized into five FCs (Fig. 2C and Table 1, S3). In FCI category, we identified 10 proteins (AtpA, ICDH, CcmM, CpcG2, PsaD, PetE, Rpe, Gap-2, HemE, and CpcB) that corresponded to 12 DAP spots (3, 6, 8, 9, 11, 18, 19, 20, 23, 31, 32, and 36) (Fig. 2C and Table 1, S3). Of these, AtpA, CpcG2, PsaD, PetE, HemE, and CpcB belong to light-dependent photosynthetic reactions, while ICDH, CcmM, Rpe, and Gap-2 are involved in carbon metabolism. AtpA, a component of the hydrophilic catalytic core of ATP synthase, was found to be down-accumulated upon LexA overexpression under unstressed conditions and increased accumulation was observed in Cd-stressed AnpAM cells compared to unstressed AnpAM cells (Fig. S1 and Table S3), but remained unchanged LexA overexpression and Cd stress was applied simultaneously (Fig. 1 and Table 1). Since *atpA* lacks an AnLexA-Box in its upstream regulatory region (Table 2) and its transcript has previously been shown not to be affected by LexA in *Anabaena* (Srivastava et al., 2022a), a decrease in its accumulation upon LexA overexpression under unstressed conditions could be an indirect effect of LexA-regulated protease causing increased degradation of AtpA. CpcG2, a rod-core linker protein for the PSI-specific peripheral antenna, CpcG2-phycobilisome, is required to stabilize the NDH-1-PSI supercomplex (Gao et al., 2016). Its accumulation significantly decreased upon LexA overexpression both in the presence and absence of Cd stress, while it did not change upon Cd stress in vector control AnpAM cells (Fig. 1, S1 and Table 1). The decrease in CpcG2 accumulation upon LexA overexpression is consistent with previous findings, which found that An*lexA*^+^ cells had a lower NDH-PSI supercomplex and consequently a lower NDH-mediated cyclic electron flow than AnpAM cells in both presence and absence of Cd stress (Srivastava et al., 2022a). Furthermore, the presence of AnLexA-box in the vicinity of its predicted −10 promoter (Table 2) suggests that LexA is involved in its regulation. PsaD is a PSI peripheral subunit II that stabilizes the binding of other PSI subunits (PsaC and PsaE) to the PSI core and hence plays a key role in PSI assembly (Minai et al., 1996). PsaD accumulation was found to be decreased in response to Cd stress and LexA overexpression individually as well as in combination (Fig. 1, S1 and Table 1), validating earlier findings of lower PSI tetramer levels in An*lexA*^+^ cells than AnpAM in both presence and absence of Cd stress (Srivastava et al., 2022a). However, transcript analysis done earlier revealed up-regulation of *psaD* gene transcripts by LexA in *Anabaena* (Srivastava et al., 2022a). Thus, the decreased accumulation of the protein in spite of increased transcription could be an effect of decreased stability or enhanced protein turnover of PsaD in the presence of LexA and/or Cd stress. PetE is a copper-containing protein that facilitates the transfer of electrons from the cytochrome *b*_6_*f* complex to PSI (Schöttler et al., 2004). PetE accumulation was reduced upon Cd stress irrespective of the presence or absence of high levels of LexA protein (Fig. 1, S1 and Table 1) indicating that it may not be under the regulation of LexA, which is further confirmed by the absence of an AnLexA-Box within 100 bases of its predicted −10 region. Thus, *petE* is a Cd-responsive gene and its decreased accumulation under Cd stress is possibly to prevent PSI over-reduction and the production of ROS. HemE is a cytosolic enzyme that catalyzes the biosynthesis of tetrapyrroles, a metal-binding cofactor found in a variety of enzymes, proteins, and pigments (Czarnecki and Grimm, 2012). Lower accumulation of HemE in response to Cd stress and LexA overexpression individually and in combination (Fig. 1, S1 and Table 1), could explain the greater decline in photosynthetic pigment content in An*lexA*^+^ cells after Cd stress compared to AnpAM cells, as previously observed (Srivastava et al., 2022a). The *hemE* gene possesses AnLexA-Box which overlaped with its predicted −10 promoter (Table 2), and LexA was found to bind to its upstream region (Fig. 3), thus confirming regulation by LexA. CpcB, the β-subunit of phycocyanin exhibited lower accumulation in response to LexA overexpression and Cd stress individually but not in combination (Fig. 1, S1 and Table 1, S3). With the presence of AnLexA-box near the predicted −10 region, CpcB has previously been proven to be directly regulated by *Anabaena* LexA (Kumar et al., 2018). CcmM is a protein necessary for assembly of carboxysomes, which concentrates carbon dioxide (CO_2_) near RubisCO and improves carbon fixation efficiency (Long et al., 2018). The accumulation of CcmM was found to be significantly reduced in response to Cd stress both in the presence and absence of LexA overexpression (Fig. 1, S1 and Table 1), presumably because the enzymes of the Calvin-Benson-Bassham (CBB) cycle are very sensitive to the Cd (Krupa et al., 1993; Singh et al., 2015). The decrease in its accumulation upon LexA overexpression under unstressed conditions (Fig. S1), with the increase in CcmM transcript level (Fig. 4B) could be explained by the significant difference in size of the protein spots (8, 9) and the actual molecular mass of these proteins (Table 1). Direct regulation of *ccmM* by LexA was confirmed by the presence of AnLexA-box close to the promoter region (Table 2), and the binding of LexA to this region (Fig. 3). ICDH catalyzes the oxidative decarboxylation of isocitrate to produce *α*-ketoglutarate and CO_2_ while reducing the NADP^+^ to NADPH in the tricarboxylic acid (TCA) cycle. It was found to be down-accumulated in response to LexA overexpression and Cd stress individually but not in combination (Fig. 1, S1 and Table 1, S3). Since NADPH provides reducing power for anabolic processes and redox equilibrium, lower ICDH accumulation upon LexA overexpression (Table S3) could be one of the causes for higher ROS levels in An*lexA*^+^ cells compared to AnpAM cells (Kumar et al., 2018). In addition, the corresponding gene has a functional AnLexA-Box in its promoter region (Table 2), implying that LexA would regulate it. Rpe and Gap-2, respectively, are involved in the pentose phosphate pathway (PPP) and glycolysis. The accumulation of Rpe and Gap-2 was found to be altered by LexA overexpression in both the absence and presence of Cd stress (Fig. 1, S1 and Table 1). The regulation of LexA on their corresponding genes has previously been confirmed by *in silico* and/or EMSA (Srivastava et al., 2022b PREPRINT). Thus, the overexpression of LexA accentuated the inhibitory effect of Cd stress on photosynthesis through modulation of expression of several genes involved in photosynthesis.

TrxA and TrxO (Alr2205), which corresponded to spots 28 and 29, respectively, were the only proteins identified in the FCII category (Fig. 2C and Table 1, S3). TrxA and TrxO are two of the seven Trx found in the genome of *Anabaena* (Ehira and Ohmori, 2012). TrxA deletion has previously been demonstrated to cause growth arrest, alter cellular redox status, and interfere with photosynthesis, carbon assimilation, and the oxidative stress response in *Synechocystis* (Mallén-Ponce et al., 2021). TrxA accumulation was found to increase exclusively in response to LexA overexpression and was unchanged in response to only Cd stress and to the combined presence of LexA overexpression and Cd stress (Fig. 1, S1 and Table 1, S3). In contrast to the reduced transcript level, TrxO accumulation was unaffected by LexA overexpression at the protein level (Fig. 4B, S1), indicating the higher stability of this protein. TrxO protein accumulation was increased in both recombinant strains in response to Cd stress in order to combat the oxidative stress caused by this heavy metal, but this increase was less pronounced in An*lexA*^+^ cells than in AnpAM cells (Fig. 1, S1 and Table 1). The presence of a functional AnLexA-Box upstream of the coding region of *trxA* and *alr2205* (Table 2) and binding of LexA protein to the upstream region of *alr2205* (Fig. 3) lead to the conclusion that differences in their accumulation pattern in response to LexA overexpression in the presence and/or absence of Cd stress are due to LexA regulation. LexA has been shown to differentially regulate several of the oxidative stress alleviation genes (Kumar et al., 2018: Srivastava et al., 2022b PREPRINT), including *trxA* and *trxO* identified in the current study, indicating an important role for LexA in maintaining the redox status of *Anabaena*.

In the FCIII category, we identified 9 proteins (DnaK2, EF-Tu, OpA, PGDH, RRF, Rpl9, Rpl10, AMT, and Rps1), which belong to 9 individual DAP spots (1, 2, 4, 7, 15, 16, 26, 33, and 38) (Fig. 2C and Table 1, S3). Of these, DnaK2 and OpA are protein folding and degradation proteins, respectively, whereas the others are involved in amino acid and protein biosynthesis. The level of DnaK2 accumulation was found to be reduced by LexA overexpression individually, but not by only Cd stress or by the combined effect of Cd and LexA overexpression (Fig. 1, S1 and Table 1, S3), concurs with the earlier reported repression of *dnaK* expression by LexA in *Anabaena* (Kumar et al., 2018). Lower availability of one of the major chaperones, DnaK would result in lower rate of refolding of misfolded proteins, which are known to be accumulated in response to environmental stresses (Dantuma and Lindsten, 2010), thus rendering the cells more susceptible to the stress. Additionally, the enhanced accumulation and transcription of OpA, a protein-degrading enzyme in response to Cd stress (Fig. 4C, S1 and Table S3), would result in targeting of the misfolded/unfolded protein for degradation. The proteins involved in amino acid and protein biosynthesis, EF-Tu, PGDH, RRF, Rpl9, Rpl10, AMT, and Rps1, were also found to be affected by LexA overexpression in either presence or absence of Cd stress or both conditions (Fig. 1, S1 and Table 1, S3), suggesting that LexA regulates their accumulation. This decrease in chaperone levels and proteins involved in amino acid and protein synthesis and the increase in protease levels could thus contribute to the observed decreased accumulation of several protein spots, which have not been shown to be directly regulated by LexA in the absence or presence of Cd stress.

In the FCIV category, we identified 3 proteins (ENR, NusA, and Rbp) that corresponded to 3 individual DAP spots (22, 30, and 39) (Fig. 2C and Table 1, S3). ENR, a key enzyme in the biosynthesis of fatty acid, showed increased accumulation in response to LexA overexpression both in the presence and absence of Cd stress but was unaffected by Cd stress alone (Fig. 1, S1 and Table 1). Regulation of LexA on the corresponding gene (*all4391*) was previously demonstrated bioinformatically (Table 2, Srivastava et al., 2022b PREPRINT). NusA is involved in the elongation and termination of transcription. It has also been demonstrated to function as a molecular chaperone and in coordinating cellular responses to DNA damage (Cohen et al., 2010; Li et al., 2013). Though the expression of *nusA* transcript increased upon LexA overexpression (Fig. 4B) and the presence of AnLexA-Box in the vicinity of its promoter region (Table 2) is suggestive of a direct regulation of *nusA* by LexA, which was further validated by an EMSA study (Fig. 3). However, the abundance of protein decreased under this condition (Table S3), suggesting either high turnover or lower stability, which needs to be investigated. On the other hand, Cd stress resulted in enhanced transcription, as well as accumulation of NusA protein (Fig. 1, 4A,C, S1 and Table 1, S3). Rbp, a post-transcriptionally active regulatory protein, was found to be down-accumulated with LexA overexpression and Cd stress alone but not combined (Fig. 1, S1 and Table 1, S3). Since *rbp* has a functional AnLexA-box in its promoter region, it is expected to be regulated by LexA (Table 2).

From FCV category, we identified 12 proteins (Alr0803, All1864, All1865, Alr0882, All2425, All7597, Alr0806, All4688, All4894, All3797, All2110, and All4050) that corresponded to 13 DAP spots (5, 10, 12, 13, 14, 17, 21, 24, 25, 27, 34, 35, and 37) (Fig. 2C and Table 1, S3). Among these, All2425 and Alr0806 are annotated in the database as unknown proteins, while the rest are annotated as hypothetical proteins. The differential accumulation of these two identified unknown proteins in response to either LexA overexpression or Cd stress or both, as well as the presence of functional AnLexA-Box upstream of these genes (Fig. 1, S1 and Table 1, 2, S3), imply direct regulation by LexA. Among the 10 identified hypothetical proteins, Alr0803, which exhibited similarity with signal transduction histidine kinase, was earlier suggested not to be regulated by LexA based on *in silico* and transcript analyses, and Alr0882, which showed similarity with universal stress protein domain-containing proteins, was earlier predicted to be regulated by LexA due to the presence of AnLexA-Box near to the −10 region (Table 2, Srivastava et al., 2022b PREPRINT). All1864 shared similarities with nitroreductases, which have been implicated in response to oxidative stress (Dabravolski, 2020). All1864 accumulation was unaffected by LexA overexpression under unstressed conditions but elevated by Cd stress at both the mRNA and protein levels in both recombinant *Anabaena* strains (Fig. 1, 4 A-C, S1 and Table 1), presumably to deal with Cd-induced oxidative damage. However, the slower increase in All1864 protein accumulation in An*lexA*^+^ compared to AnpAM (Fig. 1, S1 and Table 1) might explain the greater susceptibility to Cd stress (Kumar et al., 2018; Srivastava et al., 2022a). Similar to All1864, the accumulation of All1865, which is related to oxidoreductases, All7597, which contains MaoC_dehydratase like domain, and All4894, which is related to fasciclin 1 domain-containing proteins, was unaffected by LexA overexpression but increased by Cd stress in both recombinant strains, albeit to a lesser level in An*lexA*^+^ (Fig. 1, S1 and Table 1). Since oxidoreductases and All4894 play important roles in oxidative stress response (Rai et al., 2020; Zhang et al., 2020), their slower increase in An*lexA*^+^ cells than AnpAM cells likely made An*lexA*^+^ cells more sensitive to Cd stress (Kumar et al., 2018; Srivastava et al., 2022a). In the presence and absence of Cd stress, it was found that All4688 decreased upon LexA overexpression (Fig. 1, S1 and Table 1), but no information regarding its function is currently available in databases. Similar to All4894, All3797 also exhibited the similarity with fasciclin 1 domain-containing proteins, but was found to be considerably reduced in abundance upon LexA overexpression and Cd stress individually, as well as in combination (Fig. 1, S1 and Table 1). Higher ROS levels in An*lexA*^+^ cells than AnpAM cells are likely due to less All3797 accumulation upon LexA overexpression (Kumar et al., 2018). All2110 possessed the sequence resemblance with von Willebrand factor A-like (vWFA) domain-containing proteins, which have been involved in numerous biological functions, such as adhesion, migration, signal transduction, DNA repair, membrane transport, and heavy metal tolerance (Colombatti et al., 1993; Whelan et al., 1997). The increase in All2110 accumulation upon Cd stress in AnpAM cells (Fig. S1) might be to offset the toxic impact of Cd, whereas the reduction in All2110 accumulation upon LexA overexpression under both unstressed and Cd stressed conditions (Fig. 1, S1 and Table 1) again confirmed the reduced resistance of An*lexA*^+^ cells to Cd stress (Kumar et al., 2018; Srivastava et al., 2022a). All4050 exhibited the similarity with photosynthetic reaction center (PRC)-barrel domain-containing proteins, which are required for transferring the electrons within the cyclic electron transport pathway, as well as for the maturation of 16S rRNA (Anantharaman and Aravind, 2002). During Cd stress, All4050 accumulation increased in AnpAM (Fig. S1 and Table S3), which is expected to stimulate cyclic electron flow around PSI upon Cd-induced PSII damage (Srivastava et al., 2021a, 2022b). However, the lower level of All4050 in An*lexA*^+^ cells than in AnpAM cells in both unstressed and Cd stressed conditions (Fig. S1 and Table S3) validates the lower cyclic electron flow in An*lexA*^+^ cells (Srivastava et al., 2022a). As *all1864*, *all1865*, *all7597*, *all7597*, *all4894*, *all3797*, *all2110*, and *all4050* all have an AnLexA-box near their predicted −10 regions (Table 2), they are all likely to be regulated by LexA. Among these, the direct regulation of LexA on *all1864* gene was experimentally demonstrated in Fig. 3.

### 4.2 Overexpression of LexA modulated the expression of Cd efflux transporter genes in Anabaena

Since prokaryotes have strong resistance to heavy metals due to their multiple efflux systems (Nies, 2003), we examined the transcript levels of six previously reported Cd efflux transporters, *all7618*, *all7619*, *ycf16*, *aztA*, *alr4267*, and *secD*, in response to both combined and individual Cd stress and LexA overexpression to see if they are regulated by LexA (Fig. 4D-F). Among them, *ycf16* and *aztA* are direct ATP-dependent primary transporters for toxic compound extrusion from the cytoplasm across the periplasmic space to the outside of the cell, whereas *all7618*, *all7619*, and *alr4267* are secondary transporters that are usually energized by the electrochemical gradients established by primary active transport (Li et al., 2002; Anes et al., 2015). All of these transporters have been proven previously to confer Cd tolerance in a variety of organisms, including cyanobacteria, by effluxing Cd out of the cell (Nesler et al., 2017; Xu et al., 2018; Victoria et al., 2018; Fu et al., 2019; Liu et al., 2021; Srivastava et al., 2021b). The transcript levels of *ycf1* (*alr2493*), an ABC transporter ATP-binding protein, which is organized in the form of an operon (*alr2492*-*alr2493*-*alr2494*-*alr2495*-*alr2496*), and a single gene *aztA*, a Cd-specific P-type ATPase, increased in response to LexA overexpression under unstressed condition (Fig. 4E, Table 2). When exposed to Cd stress, the transcript level of *ycf1* increased in both strains, but to a higher extent in An*lexA*^+^ cells than in AnpAM cells, while the transcript level of *aztA* was unaffected in either strain (Fig. 4D,F), suggesting lower energy conservation in An*lexA*^+^ cells. The change in the mRNA levels of *ycf1* and *aztA* due to LexA overexpression under both unstressed and Cd stressed conditions (Fig. 4D-F), as well as the presence of AnLexA-box in the vicinity of their predicted −10 regions (Table 2), imply that LexA regulates them, which was further validated by LexA binding studies to the upstream regulatory regions (Fig. 3). The transcript levels of *all7618* and *all7619*, which are homologous to *czcA* and *czcB* of the three-component CzcCBA chemiosmotic efflux pump and are arranged as *all7619*-*all7618*-*all7617* operon, and *alr4267*, a cation or drug efflux protein that functions as *alr4267*-*alr4268*-*alr4269*, increased in response to LexA overexpression under unstressed condition (Fig. 4E and Table 2). However, when subjected to Cd stress, *all7618*, *all7619*, and *alr4267* transcripts decreased in vector control AnpAM cells (Fig. 4F), while they remained unaltered in An*lexA*^+^ cells (Fig. 4D), again indicating less energy conservation due to LexA overexpression. Furthermore, the presence of functional AnLexA-boxes in their promoter regions (Table 2) indicates that all three are regulated by LexA. LexA binding in the promoter region of *all7619* provided further evidence for the direct regulation of the genes *all7618* and *all7619* (Fig. 3). Gram-negative bacteria export most proteins from the cytoplasm into and across the plasma membrane through a large multimeric protein export complex called sec-translocase, and SecD is a component of this complex that promotes translocation efficiency (Pogliano and Beckwith, 1994). The transcript level of *secD* increased in response to both individual and combined LexA overexpression and Cd stress (Fig. 4D-F), but because it lacks functional AnLexA-box in its promoter regions (Table 2), LexA regulation on this gene may be indirect.

## Conclusions

In the current study, cytosolic proteome analysis of *Anabaena* overexpressing LexA protein in response to Cd stress, followed by RT-qPCR and EMSA validation, showed that in addition to modulating photosynthetic redox poising under Cd stress as reported previously (Srivastava et al., 2022a), LexA modulates the expression of a number of Cd stress-responsive genes and proteins related to photosynthesis, carbon metabolism, antioxidants, protein turnover, post-transcriptional modifications, Cd efflux, and a few unknown and hypothetical proteins that collectively coordinate the response of *Anabaena* to Cd stress.

## Acknowledgements

A.S. thanks to Department of Science and Technology-Innovation of Science Pursuit for Inspire Research (DST-INSPIRE, New Delhi, India) fellowship and S.B. thanks to Council of Scientific & Industrial Research (CSIR, New Delhi, India) fellowship for PhD. Y.M. acknowledges the financial support provided by the Department of Atomic Energy-Board of Research in Nuclear Sciences (DAE-BRNS, Mumbai, India) (Grant number: 37(1)/14/12/2017-BRNS/37209. He also acknowledges Institute of eminence (IOE) incentive grant, Banaras Hindu University (BHU, Varanasi, India) (R/Dev/D/IOE/Incentive/2021-22/32401). We are thankful to Prof. Ram Sagar (Department of Botany, Banaras Hindu University, Varanasi, India) for statistical analysis. We would also like to express our gratitude to the Head and the Programme Coordinator (CAS) in Botany and Interdisciplinary School of Life Science (ISLS), at Banaras Hindu University, Varanasi, India, for providing other instrumental support.

## Author Contributions

A.S., A.K., H.R. and Y.M. conceived the idea and designed the experiments. A.S., A.K., and S.B. conducted the experiments. A.S., V.S., H.R., and Y.M. analyzed the data and wrote the manuscript with the help of the remaining authors.

## Supplementary Files

**Table S1.**
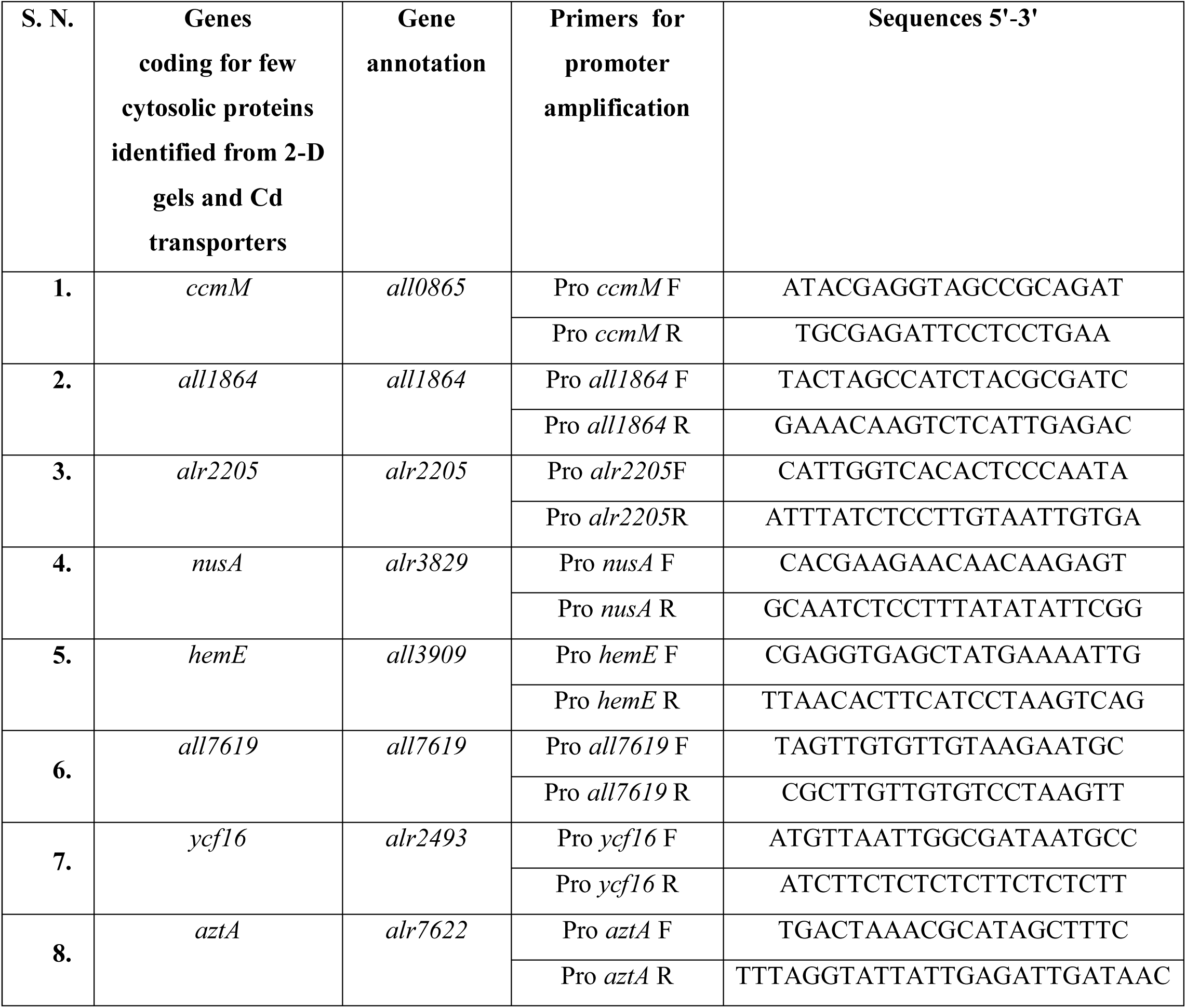
List of primers used for electrophoretic mobility shift assay (EMSA)

**Table S2.**
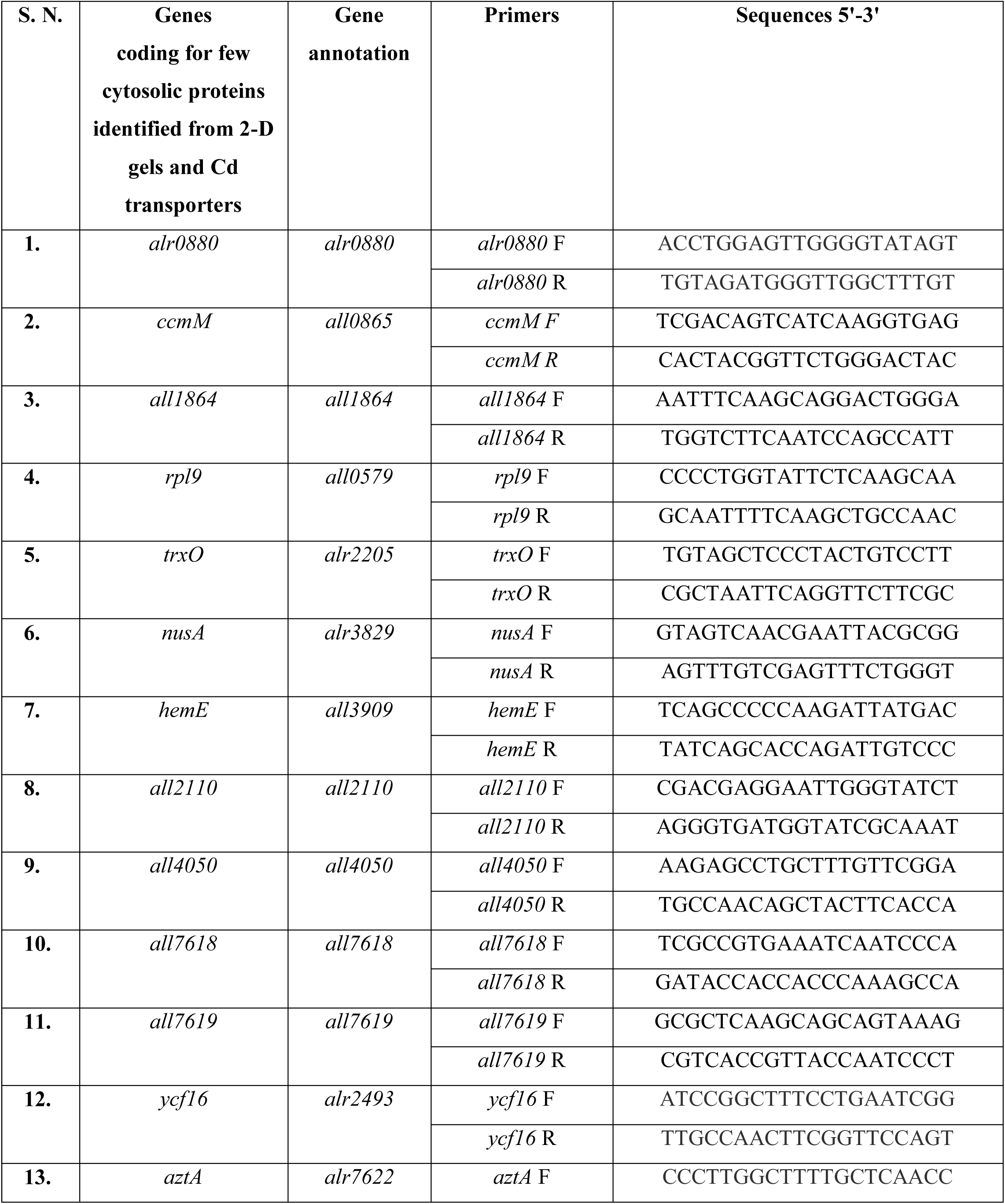
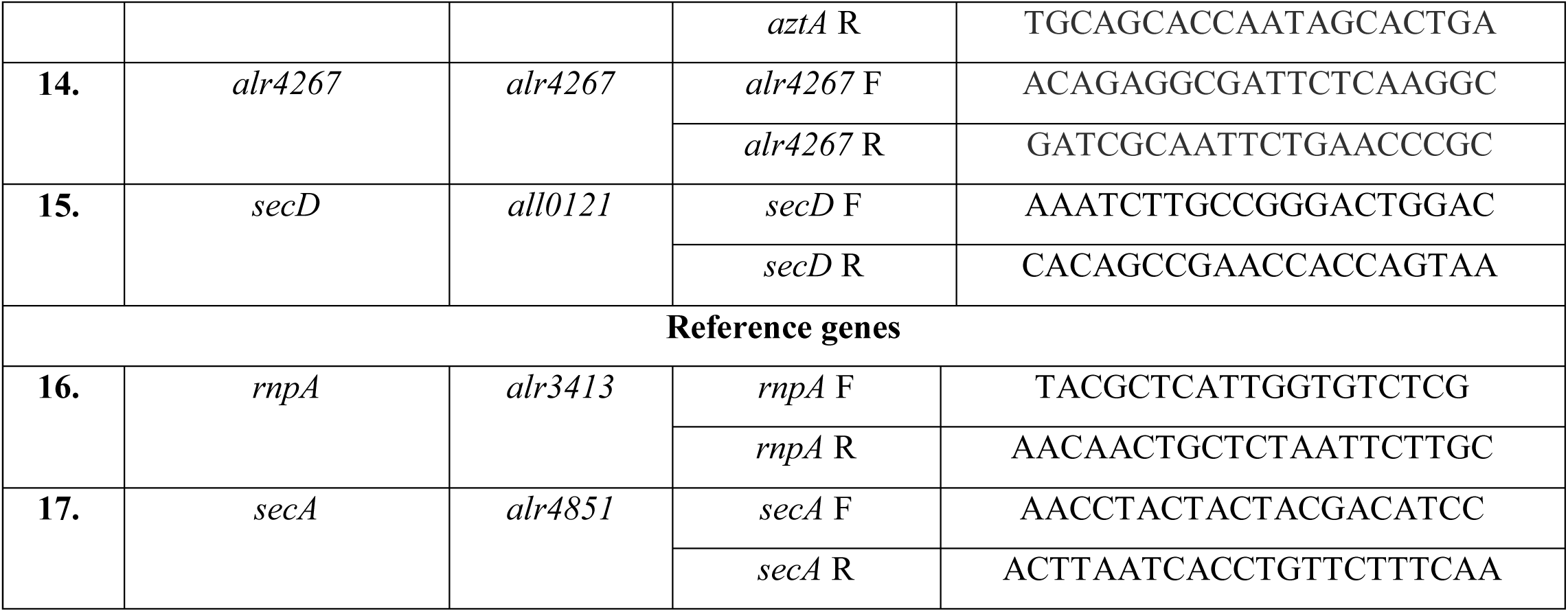
List of primers used for real time-quantitative PCR (RT-qPCR)

**Fig. S1.**
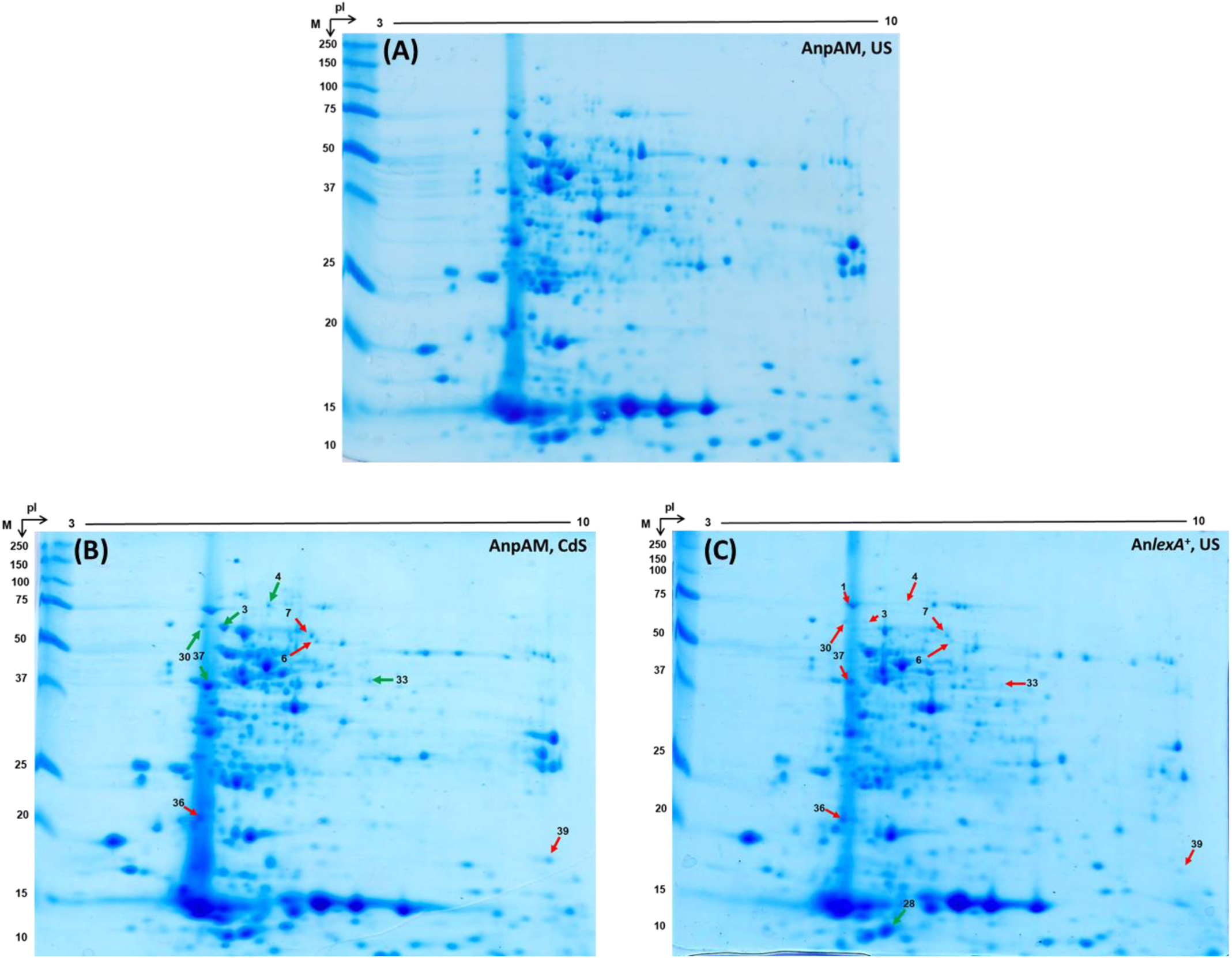
Two-dimensional (2-D) cytosolic proteome of (A and B) vector control AnpAM cells under (A) unstressed conditions, US and (B) in response to 1 d of Cd stress, CdS, and (C) unstressed LexA-overexpressing An*lexA*^+^ cells. Differentially accumulated protein (DAP) spots exclusively in response to (i) Cd stress i.e. comparison between the gels ‘A’ and ‘B’ are marked in gel B, or (ii) LexA overexpression i.e. comparison between the gels ‘A’ and ‘C’ are marked in gel C. DAP spots with increasing accumulation are shown by green arrows, whereas those with decreased accumulation are denoted by red arrows. The DAP spots, which were observed in response to the combined presence of LexA and Cd stress and shown in Fig. 1 are not included in these gels. The molecular masses (kDa) of protein size markers are displayed on the left, while the pI (3-10) is presented on the top.

**Table S3.**
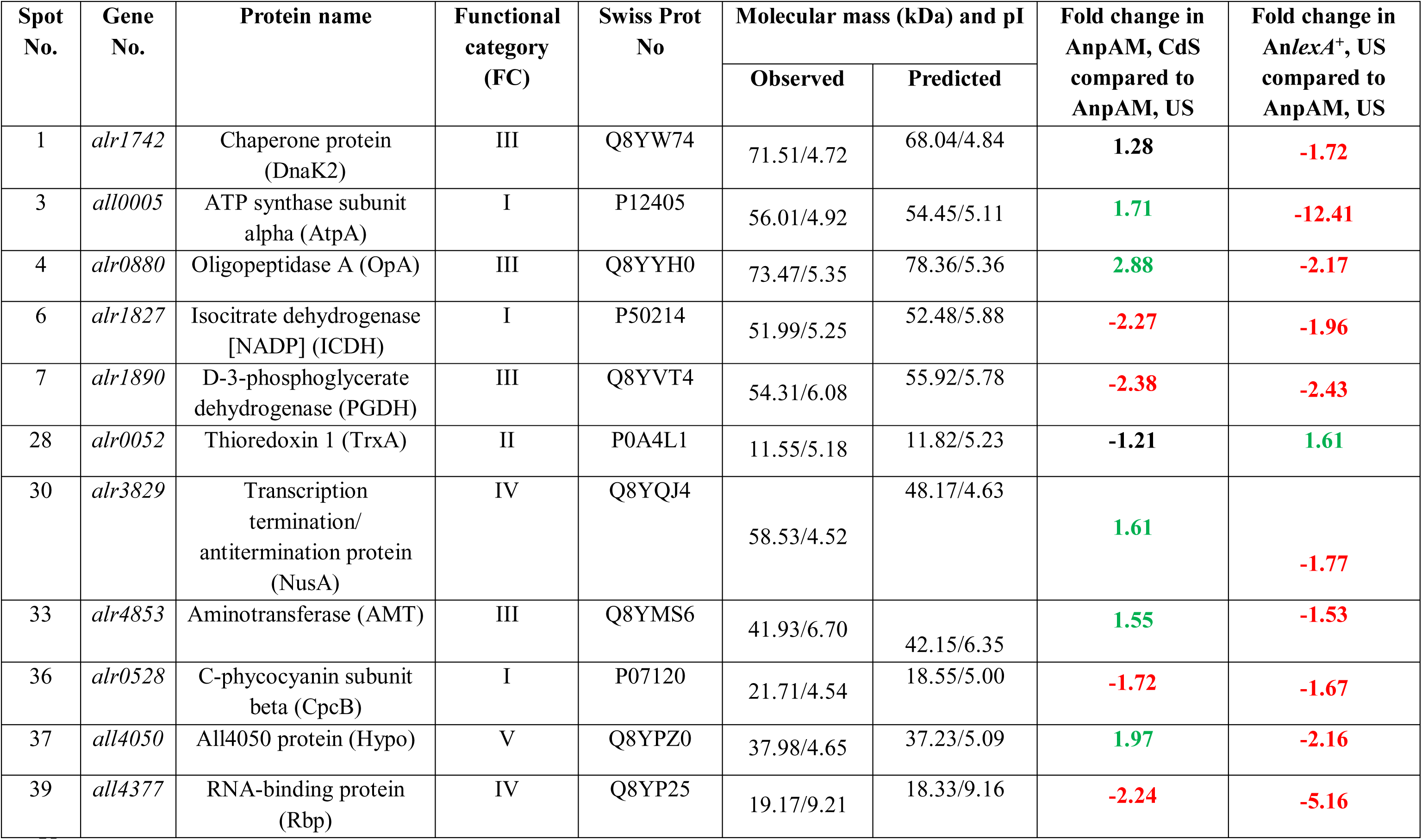
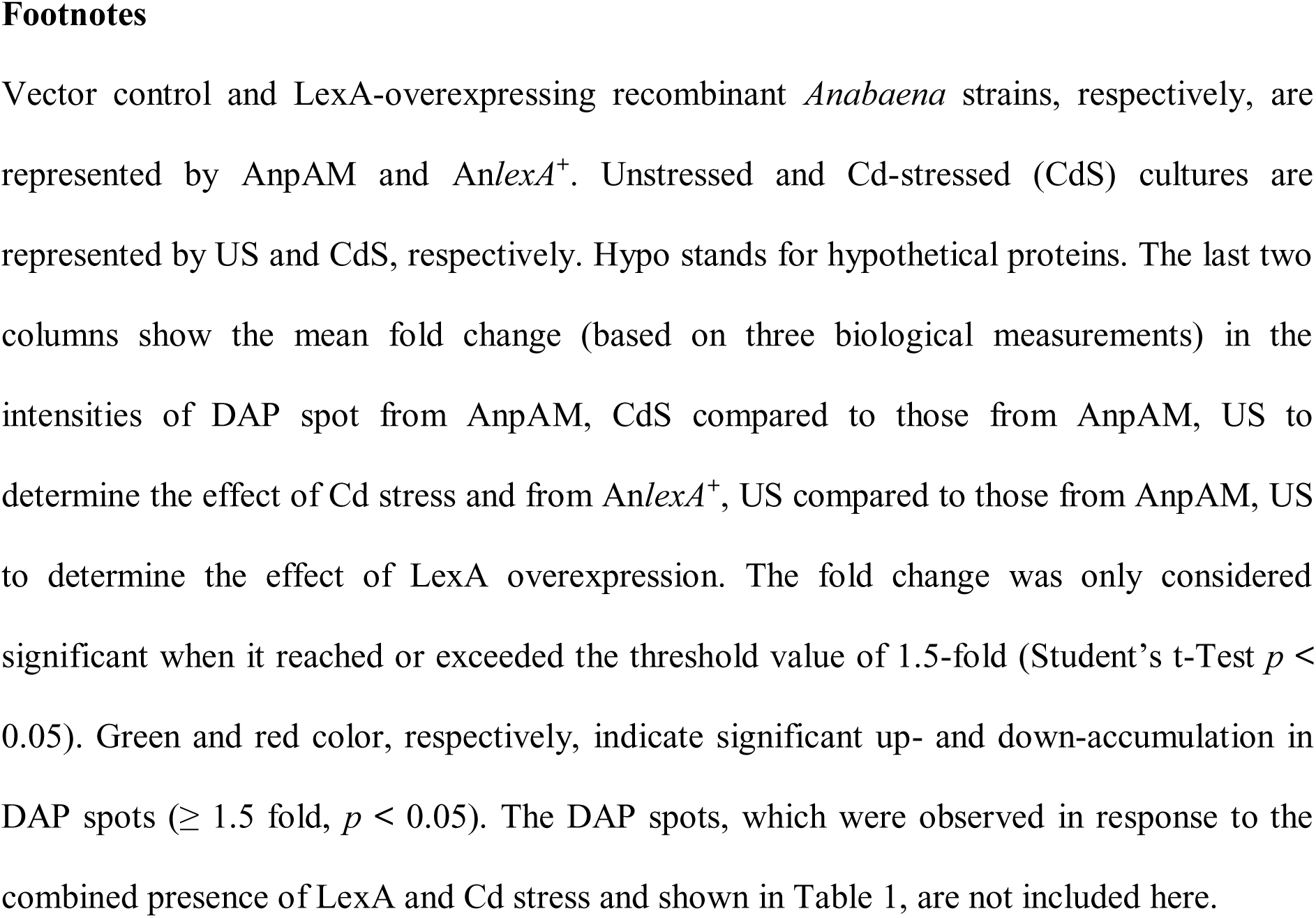
Identification details, as well as mean fold changes in the intensities of differentially accumulated protein (DAP) spots in *Anabaena* sp. PCC7120 exclusively in response to Cd stress or LexA overexpression

